# Restructuring of genomic provinces of surface ocean plankton under climate change

**DOI:** 10.1101/2020.10.20.347237

**Authors:** Paul Frémont, Marion Gehlen, Mathieu Vrac, Jade Leconte, Tom O. Delmont, Patrick Wincker, Daniele Iudicone, Olivier Jaillon

## Abstract

The impact of climate change on diversity, functioning and biogeography of marine plankton remains a major unresolved issue. Here, niche theory is applied to plankton metagenomes of 6 size fractions, from viruses to meso-zooplankton, sampled during the *Tara* Oceans expedition. Niches are used to derive plankton size-dependent structuring of the oceans south of 60°N in *climato-genomic* provinces characterized by signature genomes. By 2090, assuming the RCP8.5 high warming scenario, provinces would be reorganized over half of the considered ocean area and quasi-systematically displaced poleward. Particularly, tropical provinces would expand at the expense of temperate ones. Sea surface temperature is identified as the main driver of changes (50%) followed by phosphate (11%) and salinity (10%). Compositional shifts among key planktonic groups suggest impacts on the nitrogen and carbon cycles. Provinces are linked to estimates of carbon export fluxes which are projected to decrease on average by 4% in response to biogeographical restructuring.

---

Planktonic communities are composed of complex and heterogeneous assemblages of small animals, single-celled eukaryotes (protists), bacteria, archaea and viruses - that drift with currents. They contribute to the regulation of the Earth system through primary production via photosynthesis^1^, carbon export to the deep ocean^2, 3^ and are at the base of the food webs that sustain the whole trophic chain in the oceans^4^.

The composition of communities varies over time at a given site with daily^5^ to seasonal fluctuations^6^ following environmental variability^7^. Overlying these relatively short scale spatio-temporal variations, a more macroscale partitioning of the ocean has been revealed by different combinations of biological and physico-chemical data^8–10^, and recently documented at the resolution of community genomics^11^. The basin scale biogeographical structure has been proposed to result from a combination of multiple bio-physico-chemical processes named the seascape^7^. These processes include both abiotic and biotic interactions^12^, neutral genetic drift^13^, natural selection^14, 15^, temperature variations, nutrient supply but also advection and mixing along currents^11, 13^.

Today, knowledge of global scale plankton biogeography at the DNA level is in its infancy. We lack understanding and theoretical explanations for the emergence and maintenance of biogeographical patterns at genomic resolution. Omics data (*i.e.* the DNA/RNA sequences representative of the variety of coding and non-coding sequences of organisms) provide the appropriate resolution to track and record global biogeographical features^11^, modulation of the repertoire of expressed genes in a community in response to environmental conditions^2, 16, 17^ and eco-evolutionary processes^13–15^. Moreover, metagenomic sequencing can be consistently analyzed across plankton organisms as recently demonstrated by global expeditions^2, 18–21^. The strong links between plankton and environmental conditions suggest potentially major consequences of climate change on community composition and biogeography^22, 23^. Time series observations have highlighted recent changes in the planktonic ecosystem attributed to anthropogenic pressures, such as changes in community composition^24^ or poleward shifts of some species^25^. These changes are expected to intensify with ongoing global warming^26^ and could lead to major reorganization of plankton community composition^22^, with a potential decline in diversity^27–29^. Another major consequence of global reorganization of the seascape on biological systems would be a decrease of primary production at mid-latitudes and an increase at higher latitudes^26^.

Here we report the global structure of plankton biogeography south of 60°N based on metagenomic data using niche models and its putative modifications under climate change. First, we define environmental niches^30^, *i.e.* the envelope of environmental parameters suitable for an organism or a population, at the scale of genomic provinces across 6 organism size fractions representing major plankton groups from nano- (viruses) to meso-zooplankton (small metazoans). Then, we spatially extrapolate their niches into *climato-genomic* provinces to derive the structure of plankton biogeography for each size fraction individually and for all combined. Next, considering the same niches, we assess the spatial reorganization of these provinces under climate change by the end of the century. We study compositional changes among planktonic groups known to be critical for two major biogeochemical cycles: nitrogen with nitrogen-fixing bacteria, and carbon with phototrophs and copepods. By correlating our provinces with carbon export fluxes, these compositional changes would lead to a reduction of approximately 4% of carbon export. Finally, we quantify the relative importance of the environmental drivers explaining projected changes.

## Niche models and signature genomes from genomic provinces

We define and validate environmental niches using 4 machine learning techniques for 27 previously defined genomic provinces^11^; they correspond to 529 metagenomes for 6 size fractions (ranging from 0 to 2000 µm) sampled at 95 sites from all oceans except the Arctic (Supplementary information 1, Extended Fig 1, Supplementary Fig 1). Predictor variables of the niches are sea surface temperature (SST), salinity, dissolved silica, nitrate, phosphate and iron, plus a seasonality index of nitrate.

The signal of ocean partitioning is likely due to abundant and compact genomes whose geographical distributions closely match provinces. Within a collection of 1778 bacterial, 110 archaeal and 713 eukaryotic environmental genomes^31, 32^ characterized from *Tara* Oceans samples without cultivation, we find a total of 324 signature genomes covering all but 4 provinces, and displaying a taxonomic signal coherent with the size fractions (Fig. 1 for eukaryotes and Extended Data 2 for Bacteria and Archaea). Some of the signature genomes correspond to unexplored lineages with no cultured representatives, highlighting the knowledge gap for organisms that structure plankton biogeography and the strength of a rationale devoid of any *a priori* on reference genomes or species.

**Fig. 1.**
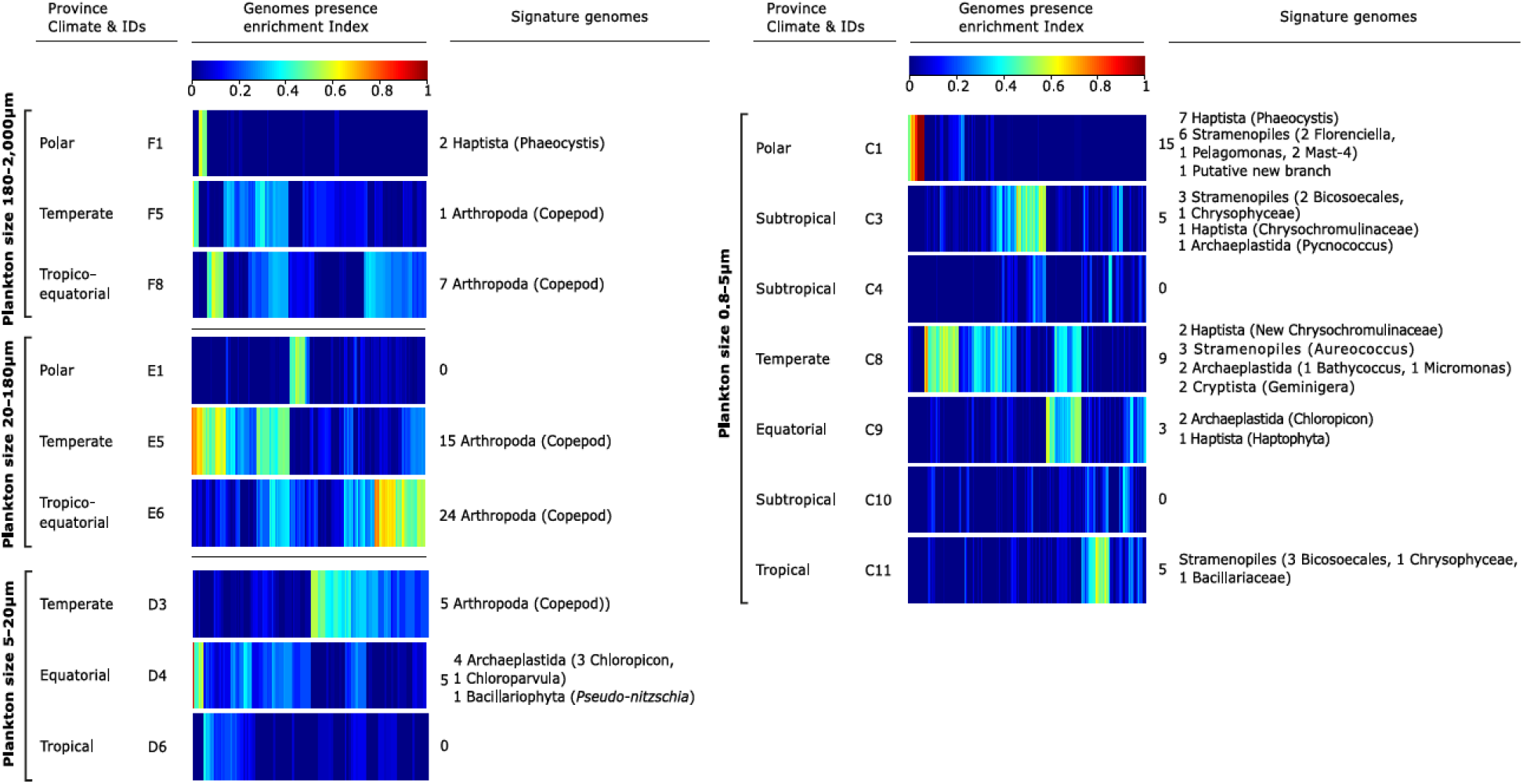
Eukaryotic signature genomes of provinces of eukaryote enriched size classes. For each plankton size class, indexes of presence enrichment (equation (1)) for 713 genomes of eukaryotic plankton^31^ in corresponding provinces are clustered and represented in a color scale. Signature genomes (see *Methods*) are found for almost all provinces, their number and taxonomies are summarized (detailed list in Supplementary Table 6).

## Structure of present day biogeography of plankton

To extrapolate the niches to a global ocean biogeography for each size fraction, we define the most probable provinces, named hereafter as *dominant* and assigned to a climatic annotation (Supplementary Table 1), on each 1°x1° resolution grid using 2006-13 WOA13 climatology^33^ (Supplementary Fig. 2 and Fig. 2).

**Fig. 2.**
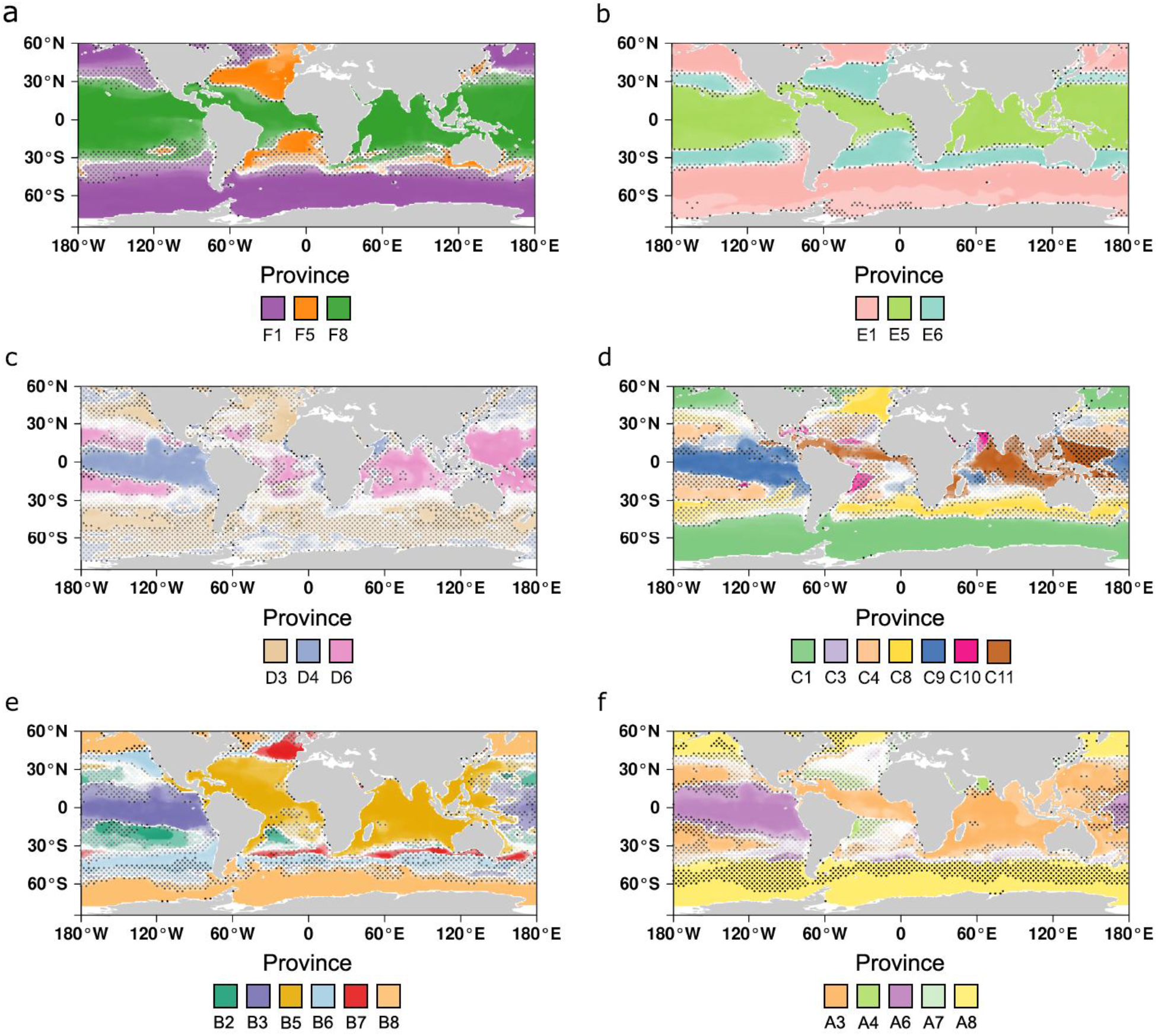
Global biogeographies of size-structured plankton provinces projected on WOA2013 dataset. (**a**) Metazoans enriched (180-2000 µm) (**b**) Small metazoans enriched (20-180 µm) (**c**) protist enriched (5-20 µm) (**d**) protist enriched (0.8-5 µm) (**e**) Bacteria enriched (0.22-3 µm) (**f**) Viruses enriched (0-0.2 µm). (**a-f**) Dotted areas represent uncertainty areas where the delta of presence probability of the dominant province and another (from the same size fraction) is inferior to 0.5. Simple biogeographies are observed in large size fractions (>20 µm) with a partitioning in three major oceanic areas: tropico-equatorial, temperate and polar. More complex geographic patterns and patchiness are observed in smaller size fractions.

In agreement with previous observations^11^, provinces of large size fractions (>20 µm) are wider and partially decoupled from those of smaller size fractions, probably due to differential responses to oceanic circulation and environmental variations, different life cycle constraints, lifestyles^7, 11^ and trophic network positions^34^. Biogeographies of small metazoans that enrich the largest size fractions (180-2000 and 20-180 µm) are broadly aligned with latitudinal bands (tropico-equatorial, temperate and (sub)-polar) dominated by a single province (Fig. 2ab). A more complex oceanic structuring emerges for the smaller size fractions (<20 µm) (Fig. 2c-f) with several provinces per large geographical region. For size fraction 0.8-5 µm enriched in small protists (Fig. 2d), distinct provinces are identified for tropical oligotrophic gyres and for the nutrient-rich equatorial upwelling region. A complex pattern of provinces, mostly latitudinal, is found for the bacteria (Fig. 2e, 0.22-3 µm) and the virus enriched size classes (Fig. 2f, 0-0.2 µm) although less clearly linked to large-scale oceanographic regions. A single province extending from temperate to polar regions emerges for size fraction 5-20 µm enriched in protists (Fig. 2c), for which fewer samples were available (Supplementary Fig. 1b-c), a likely source of bias. A consensus map combining all size fractions, built using the PHATE algorithm^35^, summarizes the main characteristics of the biogeographies described above (Supplementary information 2 and Supplementary Fig. 3).

Finally, we compare genomic biogeographies and existing ocean partitionings^8–10^. Though each of them is unique (Supplementary Fig. 6-8), common borders highlight a global latitudinal partitioning independent of the type of data (Supplementary information 3).

## Future changes in plankton biogeography structure

We assess the impacts of climate change on plankton biogeography at the end of the century following the Representative Concentration Pathway 8.5 (RCP8.5)^36^ greenhouse gas concentration trajectory. To consistently compare projections of present and future biogeographies, we use the bias-adjusted mean of 6 Earth System Model (ESM) climatologies (Supplementary Table 2, Supplementary Fig. 9). The highest warming (7.2°C) is located off the east coast of Canada in the North Atlantic while complex patterns of salinity and nutrient variations are projected in all oceans (Supplementary Fig. 10). Following this trajectory, future temperature at most sampling sites will be higher than the mean and maximum contemporary temperature within their current province (Extended Data 3).

Our projections indicate multiple large-scale changes in biogeographical structure including expansions, shrinkages and shifts in plankton organism size-dependent provinces (Fig. 3a-d, Extended Data 4, Supplementary Fig. 11-12). A change in the *dominant* province in at least one size fraction would occur over 60.1% of the ocean surface studied, ranging from 12% (20-180 µm) to 31% (0.8-5 µm) (Fig. 3, Table 1). Consistent with previous studies^22, 37^, the majority of shifts are poleward (Supplementary information 2, 4 and 5).

**Fig. 3.**
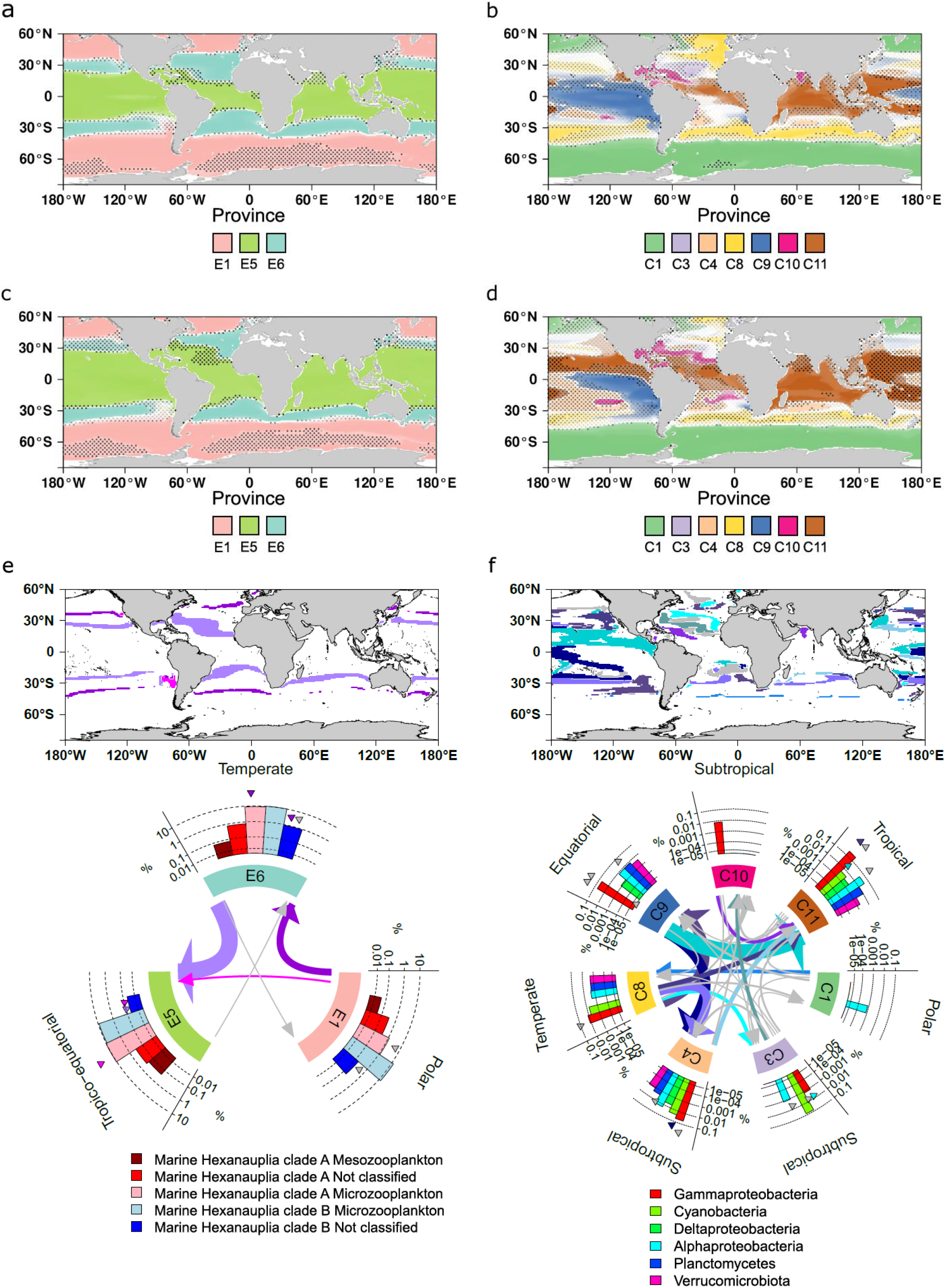
Present and future global biogeographies of the small metazoan enriched size fraction (20-180 μm) and the protist enriched size fracton (0.8-5 μm) associated compositional shifts in marine hexanauplia and diazotrohps. (**a-d**) The *dominant* province is represented at each grid point. The color transparency is the probability of presence of the province. (**e-f**) Top: Locations of *dominant* province shifts in size fraction (**e**) 20-180 μm and (**f**) 0.8-5 μm. Bottom: Summary of significant compositional shifts in (**f**) Marine Hexanauplia (copepods) or (**e**) bacterial diazotrophs: each type of transition is colored according to the one on the map or in grey (when less frequent). Barplots represent mean relative abundances based on genome abundance in the given province. Significant compositional changes in a type of genome are represented by triangles of the associated transition color.

**Table 1.** Global statistics of covered areas and province changes and transitions. From 12% to 31% of the total covered area is estimated to be replaced by a different province at the end of the century compared to present day depending on the size fraction. In total, considering all size fractions, this represents 60% of the total covered area with at least one predicted change of dominant province across the six size fractions.

| Size fraction (µm) | Present day covered area (%) | End of century covered area (%) | % area with a change of dominant province | Most frequent transition | % | 2<sup>nd</sup> most frequent transition | % |
| --- | --- | --- | --- | --- | --- | --- | --- |
| 180-2000 | 74 | 74 | 13 | temperate->tropico-equatorial (F5->F8) | 67 | polar->temperate (F1->F5) | 14 |
| 20-180 | 78 | 77 | 12 | temperate->tropico-equatorial (E6->E5) | 67 | polar->temperate (E1->E6) | 29 |
| 5-20 | 45 | 49 | 22 | temperate -> equatorial (D3->D4) | 47 | equatorial -> tropical (D4->D6) | 25 |
| 0.8-5 | 56 | 59 | 31 | equatorial -> tropical (C9->C11) | 22 | temperate -> equatorial (C8->C9) | 15 |
| 0.22-3 | 60 | 61 | 15 | temperate-> tropical (B7->B5) | 22 | polar -> temperate (B8->B6) | 16 |
| 0-0.2 | 64 | 66 | 16 | equatorial -> tropical (A6->A3) | 32 | temperate -> equatorial (A8->A6) | 31 |
| Total |  |  | 60 |  |  |  |  |

We calculate a dissimilarity index (equation (3)) at each grid point between probabilities of future and present *dominant* provinces for all size fractions combined (Fig. 4a). Large dissimilarities are obtained over northern (25° to 60°) and symmetrically southern (-25 to -60°) temperate regions (mean of 0.29 and 0.24 respectively) mostly reflecting the poleward retraction of temperate provinces (red arrows, Fig. 4a). In austral and equatorial regions, despite important environmental changes (Supplementary Fig. 10) and previously projected changes in diversity^27–29^ and biomass^38^, the contemporary provinces remain the most probable at the end of the century (mean dissimilarities of 0.18 and 0.02 respectively).

**Fig. 4.**
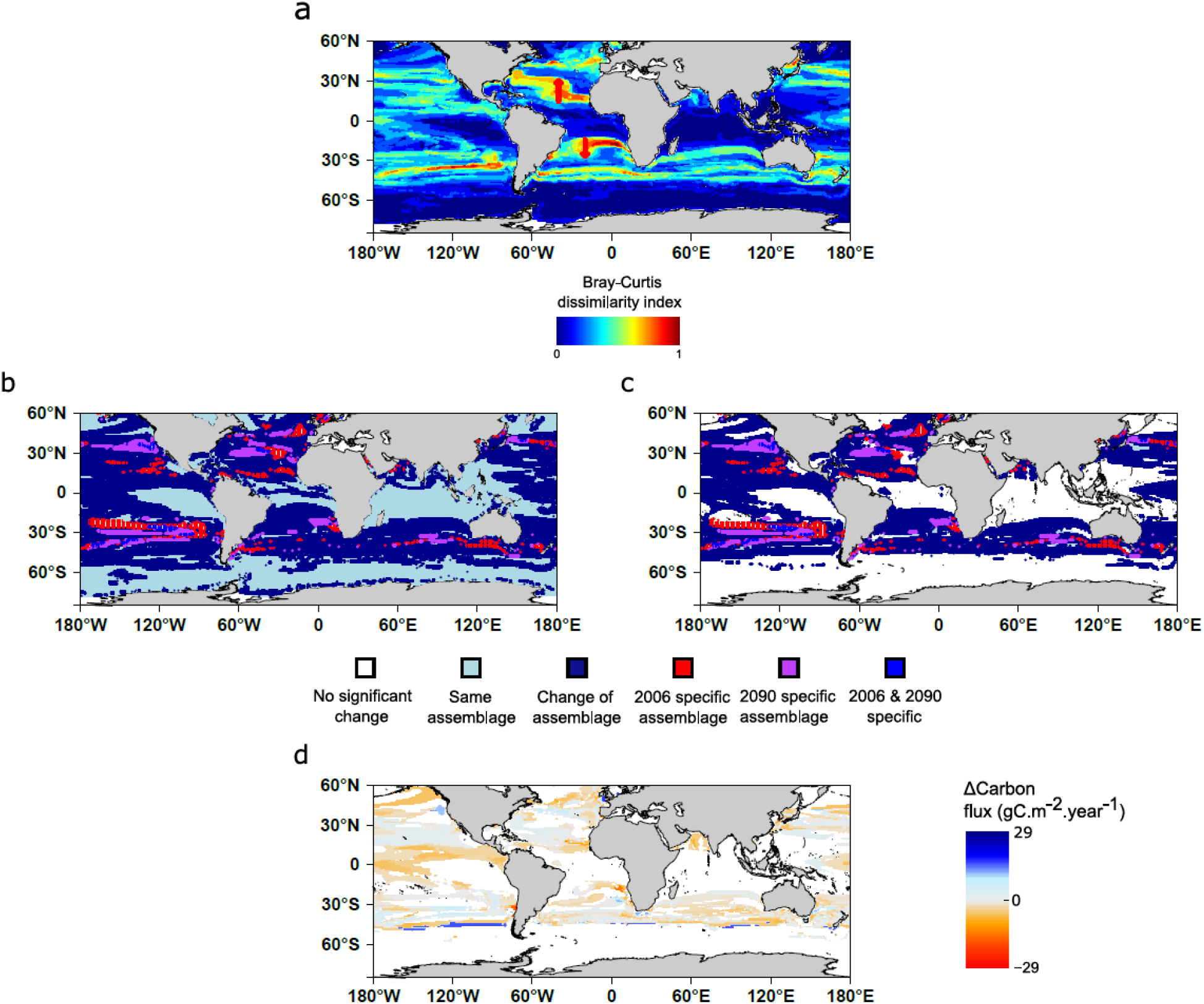
Global impact of climate change on plankton community assemblages (a-c) and carbon export (d). (**a**) Bray-Curtis dissimilarity index (equation (3)) is calculated by integrating all the *dominant* provinces presence probabilities over the six size fraction. (**b**) An assemblage is the combined projected presence of the dominant province of each size class. Assemblage reorganization is either mapped on all considered oceans (**b**) or with a criterion on the Bray-Curtis index (BC>1/6) (**c**). New assemblages are expected to appear in 2090 (purple+blue) whereas some 2006 specific assemblages are projected to disappear (red+blue). (**d**) Difference in carbon export by the end of the century based on present day mean export within each assemblage (Henson et al. 2012^3^).

To further study the future decoupling between provinces of different size fractions, we analyze the assemblages of *dominant* provinces of each size fraction. Applying two different stringent criteria, from 45.3 to 57.1% of ocean surface, mainly located in temperate regions, would be inhabited in 2090-99 by assemblages that exist elsewhere in 2006-15 (Fig. 4b versus Fig. 4c). Contemporary assemblages would disappear on 3.5 to 3.8% of the surface, and, conversely, novel assemblages, not encountered today, would cover 2.9 to 3.0% of the surface. Changes of assemblages impact key economic zones with over 41.8% to 51.8% of the surface of the main fisheries and 41.2% to 54.2% of Exclusive Economic Zones for which the future assemblage would differ from the contemporary one (Extended Data 5).

## Future changes in plankton groups important for the nitrogen and carbon cycles

To assess the potential biogeochemical impact of biogeographical restructuring, we focus on compositional changes among nitrogen-fixing bacteria (a.k.a diazotrophs), copepods and phototrophs, three groups considered important for the carbon and nitrogen cycles^39, 40^ that are well represented among the *Tara* Oceans environmental genomes. Focusing on marine areas where *dominant* provinces are projected to be replaced, we compare the present and future distribution of environmental genomes corresponding to 27 diazotrophs, 198 copepods and 231 phototrophs^31, 32^.

We project significant changes in composition for all three groups (Fig. 3e-f, Extended Data 6-9). Here we describe only changes in diazotrophs for which the genome collection most likely encompasses the whole diversity of the group (see Supplementary information 6 for changes in copepods and phototrophs).

Diazotrophy, the biotic fixation of atmospheric nitrogen, is an important process for both nitrogen and carbon cycles. It supports biological productivity in the nitrogen-limited tropical oceans^42^. Marine diazotrophs include cyanobacteria (*e.g. Trichodesmium*) described as the principal nitrogen fixers^41^ and various heterotrophic bacterial diazotrophs (HBDs) that lack cultured representatives or *in situ* imaging^32, 42^. Forty-eight of the bacterial environmental genomes are diazotrophs (8 cyanobacteria and 40 heterotrophic bacterial diazotrophs (HBDs)), encapsulating 92% of the metagenomic signal for known *nifH* genes (a reference marker for nitrogen fixation^32^) at the surface of the oceans. Twenty-seven are found in at least 5 samples and have been used for the analysis.

In size fraction 0.8-5 μm, we project significantly higher relative abundances in cyanobacteria in the tropico-equatorial regions of the Pacific Ocean (C9 to C11 and C4, Fig. 3f), which might point towards an increase in nitrogen fixation in this region as previously suggested by other models^40^. Supporting this result, we find similar significant compositional changes towards an increase in cyanobacteria in size fraction 5-20 μm (Extended Data 7c). We also project significant compositional changes for some clades of HBDs (*e.g.* increase in gammaproteobacteria: C8 to C3, Fig. 3f) though no global trend can be identified here. In the other size classes (20-180, 180-2000 and 0.22-3 μm) compositional changes are not significant (Extended Data 8). To summarize, although we cannot estimate nitrogen fixation rates using genomic data alone, genomic measurements of nitrogen-fixing cyanobacteria are in agreement with a potential increase in nitrogen fixation in the tropics, echoing results from other models^40^.

## Linking assemblages of provinces and carbon export fluxes

Carbon export refers to the processes by which organic carbon is transferred from the surface ocean to depth mainly by sinking particles that are derived from surface ocean plankton ecosystem processes^43^. Surface plankton communities are reported to be important in determining carbon export fluxes through the water column^2^. We test this hypothesis by calculating mean fluxes of carbon export that can be associated to the assemblages of our *climato-genomic* provinces from three datasets of extrapolated carbon fluxes^3, 43, 44^. We find significant differences of carbon fluxes for 46% of pairs of province assemblages (pairwise Wilcoxon test, Holm correction p<0.05). Total carbon export values are comparable to those computed from the reference data sets over the same regions (from 2.6 using Henson et al.^3^ to 6.7 PgC.year^-1^ using Eppley et al.^43^, Supplementary table 9). Our projections of the reorganization of *climate-genomic* provinces correspond to a decrease in carbon export of 4.0% on average by the end of the century (from 3.3% for Laws et al.^44^, to 4.4% for Eppley et al.^43^, Fig. 4d).

We use the Apriori algorithm^45^ to identify associations between changes in the genome composition of communities and changes in carbon export among four main latitudinal zones of the ocean: equatorial, subtropical north/south and temperate/subpolar (Extended Data 10). In temperate/subpolar regions, no significant association rules are found (Extended figure 10a). In northern and southern subtropical regions, decreases in carbon export are associated with global decreases in nano-diatoms, nano-algae and pico-algae (0.8-5 μm and 5-20) (Extended figure 10b-c). In equatorial regions, decrease in carbon export is mainly associated with decreases in nano-diatoms and nano-algae (5-20 and 0.8-5 μm) and increases in diazotrophs (0.8-5 and 5-20 μm) (Extended figure 10d). No significant rules are found for carbon export increase or changes in copepods and micro-algae.

## Drivers of plankton biogeography reorganization

We quantify the relative importance of environmental predictors (sea surface temperature, salinity, dissolved silica, phosphate, nitrate, iron and seasonality of nitrate) into niche definition and in driving future changes of the structure of plankton biogeography (equation (4)). Among these environmental properties, temperature is the first influential parameter (for 19 niches out of 27) but only at 22.6% on average (Supplementary Fig. 16a).

The relative impact of each environmental parameter is calculated^22^ for each site presenting a significant dissimilarity between 2006-15 and 2090-99 (Fig. 5a). Overall, SST would be responsible for the reorganization of the provinces at 50% followed by Phosphate (11%) and Salinity (10.3%) (Supplementary Fig. 17). Over the majority of the ocean, SST is the primary driver of the reorganization (Fig. 5a). In some regions, salinity (e.g. eastern North Atlantic) and Phosphate (e.g. equatorial region) dominate (Fig. 5a). When excluding the effect of SST, salinity and phosphate become the primary drivers of the reorganization of the provinces (Fig. 5b). The impact of SST varies across size classes with a significantly higher contribution in large size classes (>20 µm) compared to the small ones (mean of ∼73% versus ∼49%, t-test p<0.05; Fig. 5c). Though the contribution of combined nutrients to niche definition is similar for small and large size classes (mean of ∼56% versus ∼61%, Supplementary Fig. 16, Supplementary Table 3), their future projected variations have a higher relative impact on the reorganization of biogeographies of small organisms (mean of ∼39% versus ∼20%, t-test p<0.05, Supplementary Fig. 16, Supplementary Table 3). For instance, in the tropical zone, the shrinkage of the equatorial province C9 (size fraction 0.8-5 µm, Fig. 2b,d, Supplementary Fig. 18e) is driven at 24% by reduction of dissolved phosphate concentrations and at 25% by SST increase. In contrast, SST drives at 56% the shrinkage of the temperate province F5 (size fraction 180-2000 µm, Supplementary Fig. 18d). Finally, non-poleward shifts are found only within small size fractions (<20 µm) (Extended Data 4, Supplementary Fig. 11) highlighting differential responses to nutrients and SST changes between large and small organisms, the latter being enriched in phytoplankton that directly rely on nutrient supplies.

**Fig. 5.**
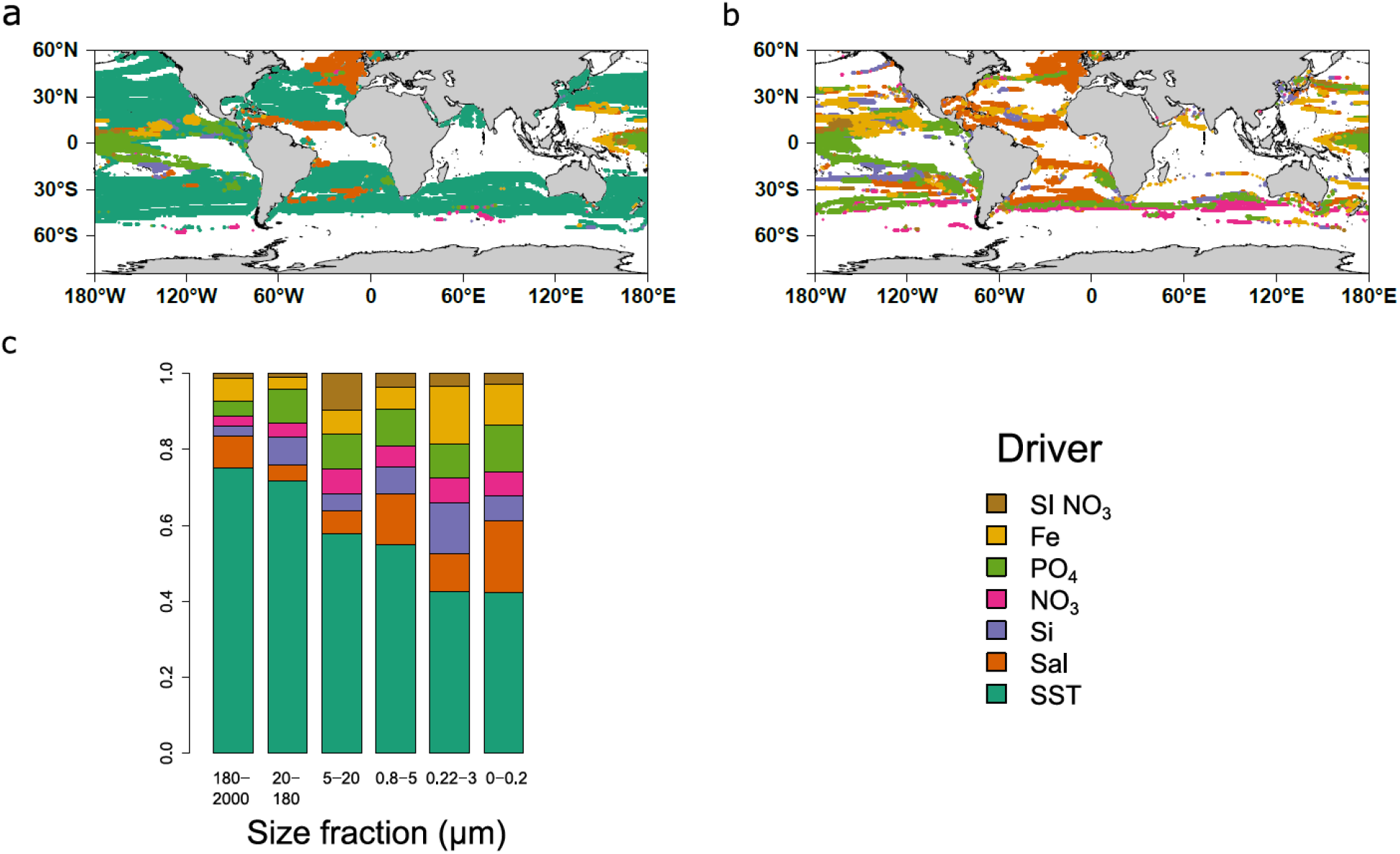
Drivers of plankton biogeographical restructuring in response to climate change. **(a)** Map of most impacting drivers on *dominant* province changes, (**b**) most impacting driver without considering temperature change and (**c**) relative importances of the drivers in the different size fractions. (**a**) Temperature appears as the top impacting driver on the majority of the ocean with a significant change of province. (**b**) Salinity and phosphate are found to be the second and third drivers of province reorganization notably at tropical and subpolar latitudes. (**c**) Temperature is found to be the most important driver for all size classes but has a more important impact in large size classes (>20 μm).

## Discussion

We propose a novel partitioning of the ocean in plankton size dependent *climato-genomic* provinces, complementing previous efforts based on other bio-physico-chemical data^9–11^. Though initially built at genomic scale, our biogeographies paradoxically reveal basin scale provinces that are larger than Longhurst’s Biogeochemical provinces^10^, and Fay and McKingley biomes^9^. These provinces are probably relatively stable across seasons as suggested by the limited effects of seasonality on the position of frontiers of BGCP provinces^10^. We propose that this apparent paradox emerges from the combination of the scale, nature and resolution of sampling. First, two proximal samples from the *Tara* Oceans expedition are separated by ∼300 km on average sampled over three years. This relatively large spatio-temporal scale overlies shorter scale compositional variations previously observed^5, 6^. Second, our estimates of plankton community dissimilarities are highly resolutive as they are computed at genomic scale with billions of small DNA fragments^11, 18^ thus smoothing out the more discrete species level signal. Together, from these combinations of processes and patterns occurring at multiple scales emerge basin scale provinces associated with coherent environmental niches and signature genomes.

These *climato-genomic* provinces are structured in broad latitudinal bands with smaller organisms (<20 μm) displaying more complex patterns and partially decoupled from larger organisms. This decoupling is the result of distinct statistical links between provinces based on organism size fractions and environmental parameters and could reflect their respective trophic modes^11, 34^.

Complex changes of the parameters defining the niches are projected under climate change leading to the reorganization of size-dependent provinces. Assuming a constant relationship to environmental drivers that define the *climato-genomic* provinces, climate change is projected to restructure them over approximately 50% of surface oceans south of 60°N by the end of the century (Fig. 4). The largest reorganization is detected in subtropical and temperate regions in agreement with other studies^29, 37^ and is accompanied by appearance and disappearance of size-fractionated provinces’ assemblages. Novel and outside of contemporary ranges environmental conditions are projected for tropico-equatorial and austral regions. While some studies extrapolate important diversity and biomass changes in these zones^27–29, 38^, here we project shifts of their boundaries and maintenance of their climatic label. The present approach does not account for putative changes in community composition or the emergence of novel niches over these regions for which novel environmental selection pressure is expected. Despite these limitations, genomic data allow us to project compositional shifts due to the reorganization of the provinces at a high phylogenetic and functional resolution.

We find significant changes in composition of diazotrophs, copepods and phototrophs including changes in size, a key trait for carbon export. These changes include increase of large copepods in subpolar regions (180-2000 μm), increase in nitrogen-fixing cyanobacteria in subtropical regions in agreement with modeling studies^40^ and complex changes in phototrophs. These changes might lead to novel prey-predator interactions *e.g.* in regions where new assemblages of communities are projected. Critically, carbon export fluxes are significantly different among *climato-genomic* province assemblages in agreement with theory^3, 26^ and previous genomic studies^2^. By the end of the century, changes in these assemblages are linked to a global decrease of the carbon pump of ∼4%, a consistent estimation compared to most models^26, 46^. This decrease is linked to decreases in nano-algae, diatoms and diazotrophs but not to copepods or micro-algae. These results are to be taken with caution especially for copepods and phototrophs as our genome database doesn’t account for the total diversity of these groups and is biased towards small and most abundant genomes. Nevertheless, the presented association of metagenomic with physico-chemical data paves the way to the integration of multiple global scale data for a better understanding of biogeochemical cycles.

Overall, our projections for the end of the century do not take into account possible future changes of major bio-physico-chemical factors such as the dynamics of community mixing, trophic interactions through transport^47^, the dynamics of the genomes^13–15^ (adaptation or acclimation) and biomass variations^38^. New sampling in current and future expeditions^48^, as well as ongoing technological improvements in bio-physico-chemical characterization of seawater samples^31, 48, 49^, will soon refine functional^16, 17^, environmental (micronutrients^50^) and phylogenetic^31^ characterization of plankton ecosystems for various biological entities (genotypes, species or communities) and spatio-temporal scales^48^. Ultimately, integrating this varied information will allow a better understanding of the conditions of emergence of ecological niches in the seascape and their response to a changing ocean.

## Supporting information

Supplementary_Table_4.genomic_provinces_WOA13_parameters

Supplementary_Table_5.genomic_provinces_niche_cross_validation

Supplementary_Table_6.genomic_provinces_signature_genomes

Supplementary_Table_7.genomic_provinces_centroid_shifts

Supplementary_Table_8.genomic_provinces_covered_areas

Supplementary_Table_9.carbon_export_fluxes_bilans

## Acknowledgments

PF was supported by a CFR doctoral fellowship and the *NEOGEN impulsion* grant from the Direction de la Recherche Fondamentale (DRF) of the CEA. This study received funding from the European Union’s Horizon 2020 Blue Growth research and innovation program under grant agreement No 862923 (project AtlantECO). This study benefited from the ESPRI computing and data centre (https://mesocentre.ipsl.fr) which is supported by CNRS, Sorbonne Université, Ecole Polytechnique and CNES as well as through national and international grants. Funding from the French state aid managed by the ANR under the “Investissements d’avenir” programme with the reference ANR-11-IDEX-0004-17-EURE-0006 » is also acknowledged. We thank Tilla Roy for preparation of the climatic data, Stephanie Henson for providing carbon export data, LAGE (Laboratoire d’Analyses Génomiques des Eucaryotes, CEA) members for stimulating discussions on this project, Mahendra Mariadassou, Sakina Dorothée Ayata and Bruno Hay Mele for discussions on statistics and climate envelope models, Laurent Bopp for initial discussions on this project and on climate models and Noan Le Bescot (TernogDesign) for help with the figures. We thank all members of the *Tara* Oceans consortium for maintaining a creative environment and for their constructive criticism. *Tara* Oceans would not exist without the *Tara* Ocean Foundation and the continuous support of 23 institutes (https://oceans.taraexpeditions.org/).

This article is contribution number XX of *Tara* Oceans.

## Competing interests

The authors declare no competing interests.

## Author contributions

PF, OJ and MG conceived the study. MV wrote the bias correction algorithm. PF computed the results, compiled and analyzed the data. PF wrote the initial draft of the paper. JL, OJ and MG conducted a preliminary study. PF, OJ, MG, MV, TD, DI, and PW discussed the results and contributed to write the paper.

## Online content

All data and codes used are available at http://www.genoscope.cns.fr/tara/SourceCodes/NCLIM-20102618_codes.tar.gz

## Materials and methods

### Genomic provinces of plankton

Environmental niches are computed for trans-kingdom plankton genomic provinces from Richter et al.^11^. They consist of the clustering of metagenomic dissimilarity matrices (based on the amount of DNA k-mers shared between pairs of samples) from 6 available size fractions with sufficient metagenomic data from the *Tara* Oceans dataset. The six size fractions (0-0.2, 0.22-3, 0.8-5, 5-20, 20-180 and 180-2000 μm) represent major plankton groups. Two large size classes (180-2000 µm and 20-180 µm) are enriched in zooplankton dominated by arthropods (mainly copepods) and cnidarians. Size classes 5-20 µm and 0.8-5 µm are enriched in smaller eukaryotic algae, such as dynophytes (5-20 µm), pelagophytes and haptophytes (0.8-5 µm). Finally, size classes 0.22-3 µm and 0-0.2 µm are respectively enriched in bacteria and viruses. Within each size fraction (from large to small), there are respectively 8, 8, 11, 6, 6 and 8 (48 in total) provinces defined in Richter et al.^11^ formed by *Tara* Oceans metagenomes (644 metagenomes sampled either at the surface (SUR) or at the Deep Chlorophyll Maximum (DCM) across 102 sites). The clustering of individual size fractions is independent.

### Genome signature of the provinces

We analyzed the distribution of 713 eukaryotic and 1888 prokaryotic genomes^31, 32^ within the genomic provinces. These genomes are Metagenome-Assembled Genomes (MAGs) obtained from *Tara* Oceans metagenomes. For each size class, we select MAGs that are present (according to a criteria defined in Delmont et al.^31^) in at least 5 samples. We computed an index of presence enrichment of MAGs within provinces as the Jaccard index^51^, defined as follows:

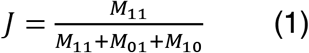

*M*_11_ is the number of samples where the MAG is present and matches a sample of the province. *M*_01 and_ *M*_10_ are respectively the number of samples where the MAG is not present in a sample of the province and inversely. A MAG is considered to be signature of a province if the Jaccard index is superior to 0.5 with this province and inferior to 0.1 for all other provinces of the given size class (Fig. 1 and Extended Data 2).

### World Ocean Atlas data

Physicochemical parameters proposed to have an impact on plankton genomic provinces^11^ are used to define environmental niches: sea surface temperature (SST), salinity (Sal); dissolved silica (Si), nitrate (NO_3_), phosphate (PO_4_), iron (Fe), and a seasonality index of nitrate (SI NO_3_). With the exception of Fe and SI NO_3_, these parameters are extracted from the gridded World Ocean Atlas 2013 (WOA13)^33^. Climatological Fe fields are provided by the biogeochemical model PISCES-v2^52^. The seasonality index of nitrate is defined as the range of nitrate concentrations in one grid cell divided by the maximum range encountered in WOA13 at the *Tara* Oceans sampling stations. All parameters are co-located with the corresponding stations and extracted at the month corresponding to the *Tara* Oceans sampling. To compensate for missing physicochemical samples in the *Tara* Oceans *in situ* data set, climatological data (WOA) are preferred. The correlation between *in situ* samples and corresponding values extracted from WOA are high (r^2^: SST: 0.96, Sal: 0.83, Si: 0.97, NO_3_: 0.83, PO_4_: 0.89). In the absence of corresponding WOA data, a search is done within 2° around the sampling location and values found within this square are averaged.

Nutrients, such as NO_3_ and PO_4_, display a strong collinearity when averaged over the global ocean (correlation of 0.95 in WOA13) which could complicate disentangling their respective contributions to niche definition. However, observations and experimental data allow identification of limiting nutrients at regional scale characterized by specific plankton communities^53^. The projection of niches into future climate would yield spurious results if the present-day collinearity is not maintained^54^ but there is up to now no evidence for large scale changes in global nutrient stoichiometry^55^.

### Earth System Models and bias correction

Outputs from 6 Earth System Models (ESM) (Supplementary Table 2) are used to project environmental niches under greenhouse gas concentration trajectory RCP8.5^36^. Environmental drivers are extracted south of 60° north for present day (2006-2015) and end of century (2090-2099) conditions for each model and the multi-model mean is computed. A bias correction method, the Cumulative Distribution Function transform, CDFt^56^, is applied to adjust the distributions of SST, Sal, Si, NO_3_ and PO_4_ of the multi-model mean to the WOA database. CDFt is based on a quantile mapping (QM) approach to reduce the bias between modeled and observed data, while accounting for climate change. Therefore, CDFt does not rely on the stationarity hypothesis and present and future distributions can be different. CDFt is applied on the global fields of the mean model simulations. By construction, CDFt preserves the ranks of the simulations to be corrected. Thus, the spatial structures of the model fields are preserved.

### Environmental niche models: training, validation and projections

Provinces with similar metagenomic content are retrieved from Richter et al.^11^. From a total of 48 initial provinces, 10 provinces are removed either because they are represented by too few samples (7 out of 10) or they are found in environments not resolved by ESMs (*e.g.* lagoons of Pacific Ocean islands, 3 out of 10). This narrows down the number of samples from 644 to 595 metagenomes. Four machine learning methods are applied to compute environmental niches for each of the 38 provinces: Gradient Boosting Machine (gbm)^57^, Random Forest (rf)^58^, single hidden layer Neural Networks (nn)^59^ and Generalized Additive Models (gam)^60^. Hyper parameters of each technique (except gam) are optimized. These are (1) for gbm, the interaction depth (1, 3 and 5), learning rate (0.01, 0.001) and the minimum number of observations in a tree node (1 to 10); (2) for rf, the number of trees (100 to 900 with step 200 and 1000 to 9000 with step 2000) and the number of parameters used for each tree (1 to 7); (3) for nn, the number of neurons of the network (1 to 10) and the decay (1.10^-4^ to 9.10^-4^ and 1.10^-5^ to 9.10^-5^). For gam the number of splines is set to 3, respectively 2 only when not enough points are available (for fraction 0-0.2, 65 points). R packages gbm (2.1.3), randomForest (4.6.14), mgcv (1.8.16) and nnet (7.3.12) are used for gbm, rf, nn and gam models.

To define the best combination of hyper parameters for each model, we perform random cross-validation by training the model on 85% of the dataset randomly sampled and by calculating the Area Under the Curve^61^ (AUC) on the 15% remaining points of the dataset. This process is repeated over 30 random subsets of the entire training set for each combination of hyperparameters. We work in a presence/absence framework *i.e.* for each province separately the dataset consists of the variable to predict (“presence” or not of the province in the sample) and the predictors consisting of the environmental variables for each sample. A fixed probability threshold of 0.5 for presence/absence detection is used to calculate the AUC. Fixing the probability threshold allows optimization of all models according to this threshold so that the *dominant* province has a reasonably high probability of presence (at least in regions with similar environmental parameters to the training dataset) and for the four types of statistical models we use. The best combination of hyper parameters is the one for which the mean AUC over the cross-validation is the highest. A model is considered valid if at least 3 out of the 4 techniques have a mean AUC superior to 0.65, which is the case for 27 out of the 38 provinces (Supplementary Fig. 1a). A climatic annotation is given to the 27 validated niches (Supplementary Table 2). Final models are trained on the full dataset and only the techniques that have a mean AUC higher than 0.65 are considered to make the projections. The vast majority (23) of the 27 validated niches is validated by all four models and 4 by only 3 models. Relative influences of each parameter in defining environmental niches are calculated using the feature_importance function from the DALEX R package^62^ for all four statistical methods (Supplementary Fig. 16a). To evaluate the consistency and coherence of environmental niche models, we first make global projections on the 2006-13 WOA2013 climatology. Projections are consistent with sampling regions for provinces encompassing vast oceanic areas. For example, the genomic province sampled in temperate Atlantic regions of size fraction 180-2000 µm is projected to be present in the north and south temperate Atlantic but also other temperate regions (Supplementary Fig. 2). For model training and projections, physicochemical variables are scaled to have a mean of 0 and a variance of 1 (using means and standard deviations of the training data). This standardization procedure allows for better performance of nn models. Finally, as statistical models often disagree on projection sets whereas they give similar predictions on the training set (Supplementary Fig. 4, 5), we use the ensemble model approach for global-scale projections of provinces^63^ *i.e.* the mean projections of the validated machine learning techniques.

### Combined size class provinces and ocean partitioning comparisons

To combine all size classes’ provinces, we use the PHATE algorithm^35, 64^ from the R package phateR. This algorithm allows visualization of high dimensional data in the requested number of dimensions while best preserving the global data structure^64^. We choose to train PHATE separately on WOA13 projections and present day and end of century projections including presence probabilities of *non dominant* provinces. We use 3 dimensions and set hyper parameter k-nearest neighbors (knn) and decay respectively to 1000 for WOA13 and 2000 for model data as in this case there are twice as many points. The hyper parameter knn reflects the degree to which the mapping of PHATE from high to low dimensionality should respect the global features of the data. We argue that 1000 and 2000 are good choices as it will be sufficient to have a highly connected graph, conserve global structure, allow visualization of structures of the size of the provinces (mean number of points in a province: 4867) and have a reasonable computational time. Decay is set to 20 in both cases. Then we cluster the resulting distance matrix using the k-medoïds algorithm^65^ and the silhouette average width criteria^66^ is used as an indicator of good fit. The silhouette criterion is maximal for 2, 3 and 4 clusters and 2 peaks are found at 7 and 14 clusters (not shown). We choose to present the 4 and 7 cluster geographical patterns as they seem more relevant with respect to the resolutions of our environmental datasets (WOA13 and climate models). We compare the three polar clusters of the 7 cluster geographical patterns with Antarctic Circumpolar Currents fronts^67^ by overlying them on the map (black lines Supplementary Fig. 3b).

To visualize the global biogeography structure, the resulting 3 vectors of PHATE are plotted using an RGB color code. Each coordinate of each vector is respectively assigned to a given degree of color component between 0 and 255 (8 bits red, green or blue) using the following formula (Supplementary Figs. 3, 13):

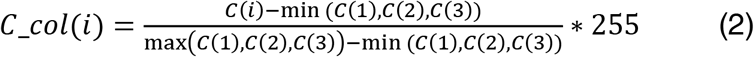

*C*(*i*) is the ith component of the PHATE axes. Respectively, components 1, 2 and 3 are assigned to red, green and blue.

To compare the six size fraction provinces, the combined size class with existing biogeochemical partitions of the oceans^9,10^ and with each other, we use the adjusted rand index^68^ (Supplementary Fig. 6-8) and overlay their masks above our partitions.

### Centroids and migration shifts

The centroid of each province is defined as the average latitude and longitude for which the probability of presence is superior to 0.5 and weighted by both the probability of presence at each grid point and the grid cell area. The migration shift is calculated as the distance between the present day and the end of the century centroids considering the earth as a perfect sphere of radius 6371 km. For consistency (*i.e.* avoid long distance aberrant shifts), it is only calculated for provinces with an area of dominance larger than 10^6^ km^2^ in the given basin.

### Bray-Curtis dissimilarity index

Climate change impact on global projections is calculated at each grid point as the Bray-Curtis dissimilarity index^69, 70^ defined as follows:

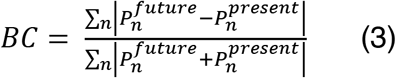

Where (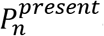 and 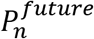) are respectively the probability of presence of the province n in present day and at the end of the century. Only the probabilities of *dominant* provinces are non-null and all others are set to zero. The mask of main fisheries^71^ (chosen as the first 4 deciles) and Exclusive Economical Zones^72^ is overlaid on the Bray-Curtis map.

### Change in province assemblages

A province assemblage is defined as the assemblage of *dominant* provinces of each size fraction at a given grid point of the considered ocean. We consider two criteria of change in province assemblage between present day and end of the century conditions. The first one, more straightforward and less stringent, considers that a province assemblage occurs when a change of *dominant* province is found in at least one size fraction. In a more stringent way, a change of assemblage is considered significant for 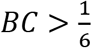 (previous section). This threshold corresponds to an idealized case where each *dominant* province has a probability of one and a change of *dominant* province is found in only one size fraction. For example, the *dominant* province assemblage goes from vector (F5,E6,D3,C8,B7,A7) (with the size fractions in decreasing order) corresponding to all temperate provinces to vector (F8,E6,D3,C8,B7,A7). This example corresponds to the replacement of the temperate province of size fraction 180-2000 µm (F5) by the tropico-equatorial province (F8). This criterion allows us to discard assemblage changes for which the changes in probability of presence of *dominant* provinces are very low.

### Composition of provinces in bacterial diazotrophs, marine copepods and phototrophs

We characterize the composition in Marine Hexanauplia (copepods), marine diazotrophs and phototrophs of provinces by considering the mean relative abundances of groups of MAGs^31, 32^ characterized taxonomically and by size (for copepods and phototrophs) in each province for all size fractions except the viral one and for size fractions >5 μm (for Marine Hexanauplia). Marine Hexanauplia are annotated taxonomically (either as belonging to clade A (109 MAGs) or clade B (105 MAGs)) from which 198 are found in at least 5 samples that we use (98 clade A and 100 clade B). To attribute a preferential size class to these MAGs, mean relative abundances over all sites of each of them are compared across size fraction 180-2000, 20-180 and 5-20 μm using Welch ANOVA^73^ (p-value<0.05). When the test is significant the MAG is annotated to its preferential size class (Mesozooplankton: 180-2000 μm, Microzooplankton: 20-180 or 5-20 μm). When the test is not significant, the MAG is annotated as unclassified. Then for each group of MAG (defined by the preferential size class plus the clade), the mean of the sum over the MAGs from this group is calculated in each province to characterize the province. The same procedure is applied for 27 out of 48 bacterial diazotrophs (present in at least 5 samples) from the prokaryotic MAG collection^32^ distinguishing groups at the phylum level (Gammaproteobacteria n=8, Cyanobacteria n=8, Deltaproteobacteria n=2, Alphaproteobacteria n=4, Planctomycetes n=3, Verrucomicrobiota n=2) and for 231 phototrophs (present in at least 5 samples) distinguishing algae (n=172), diatoms (Bacillariophyta, n=11), and cyanobacteria (n=48) and their preferential size class (Pico: 0.22-3 μm, Nano: 0.8-5 μm and 5-20 μm and Micro: 20-180 μm and 180-2000 μm). Finally, significant differential composition between provinces are annotated using Holm corrected^74^ pairwise Mann-Whitney U test^75^ (p<0.05 for significance).

### Estimation of carbon export fluxes in assemblages of provinces

We use three sets of carbon export data at 100 m depth (extrapolated data)^3, 43, 44^ to estimate an average particulate organic carbon (POC) flux for each assemblage of *climato-genomic* provinces. These extrapolations are based on estimates of carbon export efficiency from *in situ* measurements either using the *POC/*^234^*Th* radioisotope ratio^3^ or the *f-ratio* technique^43, 44^. They both derive a relation between carbon export efficiency and SST. Then to obtain extrapolations of carbon export, these ratios are multiplied by estimates of primary production from satellite estimates of sea surface chlorophyll concentrations. For each dataset, we associate each extrapolation of POC fluxes to the assemblage of present day provinces from the nearest point of the flux estimate. An average flux is associated with an assemblage if at least 3 data points are associated with that assemblage. Each grid point is associated to a given assemblage and is assigned to its average flux. Thus, we calculate carbon export estimates where an assemblage from the set of present day assemblages can be projected in the future. Next we estimate carbon export for current and projected end-of-century assemblages. Similar conclusions are reached using median values of carbon export. We verified that using whole assemblages as predictors of carbon export is more robust than using single size fraction dominant provinces (ANOVA^76^, Akaike Information Criterion^77^ (AIC)).

### Linking changes in genome composition of communities to changes in carbon export

We use the Apriori algorithm^45^ to identify associations between changes in community composition and changes in carbon export. From the relative abundance values of phototroph, diazotroph and copepod MAGs^31, 32^, we consider only significant changes in their composition among communities by a pairwise Wilcoxon test (Holm correction p<0.05). We also consider only phototrophic and copepod MAGs for which a preferred size class was found. The Apriori algorithm determines association rules between sets of ‘*transactions*’, here community changes with carbon export. The algorithm identifies sufficiently frequent associations between transactions and computes a lift as follow:

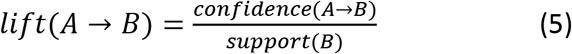

Where *confidence* (*A* → *B*) is the number of transaction where both A and B occur divided by the number of *transactions* where A occurs, and *support*(*B*) is the percentage of transactions where B occurs in the whole dataset. The lift corresponds to the increase factor in likeliness that B occurs when A occurs. The Apriori algorithm is launched for four latitudinal zones of the ocean: equatorial, subtropical (north and south) and temperate/subpolar zones. Only the directionality of changes is considered (increase/decrease in either carbon flux or a type of organism). If rules overlap perfectly, only largest association rules are kept (maximum rule length is set to 7). Only association rules pointing towards changes in carbon fluxes are kept (B=change in carbon export in equation (5)). Minimum support is set to 0.05.

### Driver analysis

To assess the relative importance of each driver in province changes, the methodology from Barton et al.^22^ is adopted. For a set N of n provinces (individual provinces or all provinces together), the probability of presence of each province is recalculated for present day conditions except for driver d (from the set of drivers D) for which the end of the century condition is used (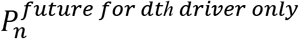). The set of driver D can be either all drivers (Fig. 5a,c) or all drivers except SST (Fig. 5b). The relative importance of driver d at a given grid point for the set of N of provinces is computed as follows:

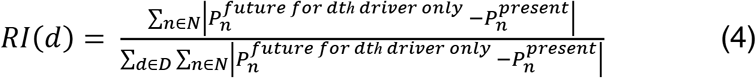

*RI*(*d*) is computed at grid cells where 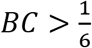 and calculated with either the set of all drivers (Fig. 5a,c) or all drivers except SST (Fig. 5b). When RI(d) is calculated for individual provinces (Fig. 5c and Supplementary Fig. 18d,e), it is computed only at grid cells where 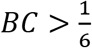 and the concerned province is either *dominant* in present day and/or end of century conditions.

---

**Extended Data 1.**
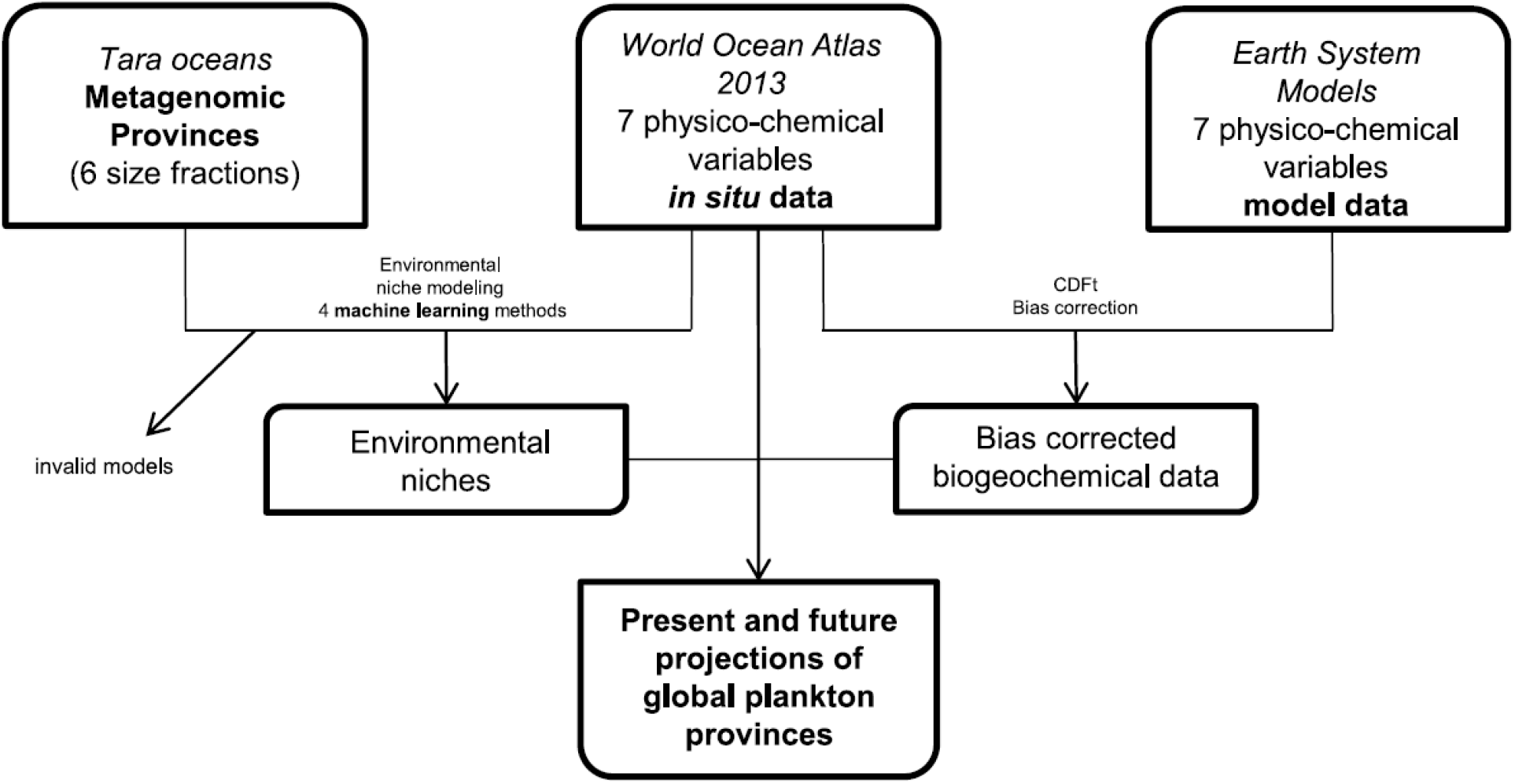
Study pipeline. Metagenomic data from the 2009-2013 *Tara Oceans* expedition and *in situ* measurements of physicochemical variables (*World Ocean Atlas 2013*, WOA13)^33^ are combined to define environmental niches at the plankton community level across 6 size fractions. Bias corrected outputs from a mean model of 6 Earth System Models (Supplementary Table 1) and WOA13 data are then used to project global plankton provinces for present day and end of the century conditions under a high warming scenario (RCP8.5)^36^. Variables are Sea Surface Temperature (SST), Salinity (Sal), Dissolved silica (Si), Nitrate (NO_3_), Phosphate (PO4), Iron (Fe) and a seasonality index of nitrate (SI NO_3_).

**Extended Data 2.**
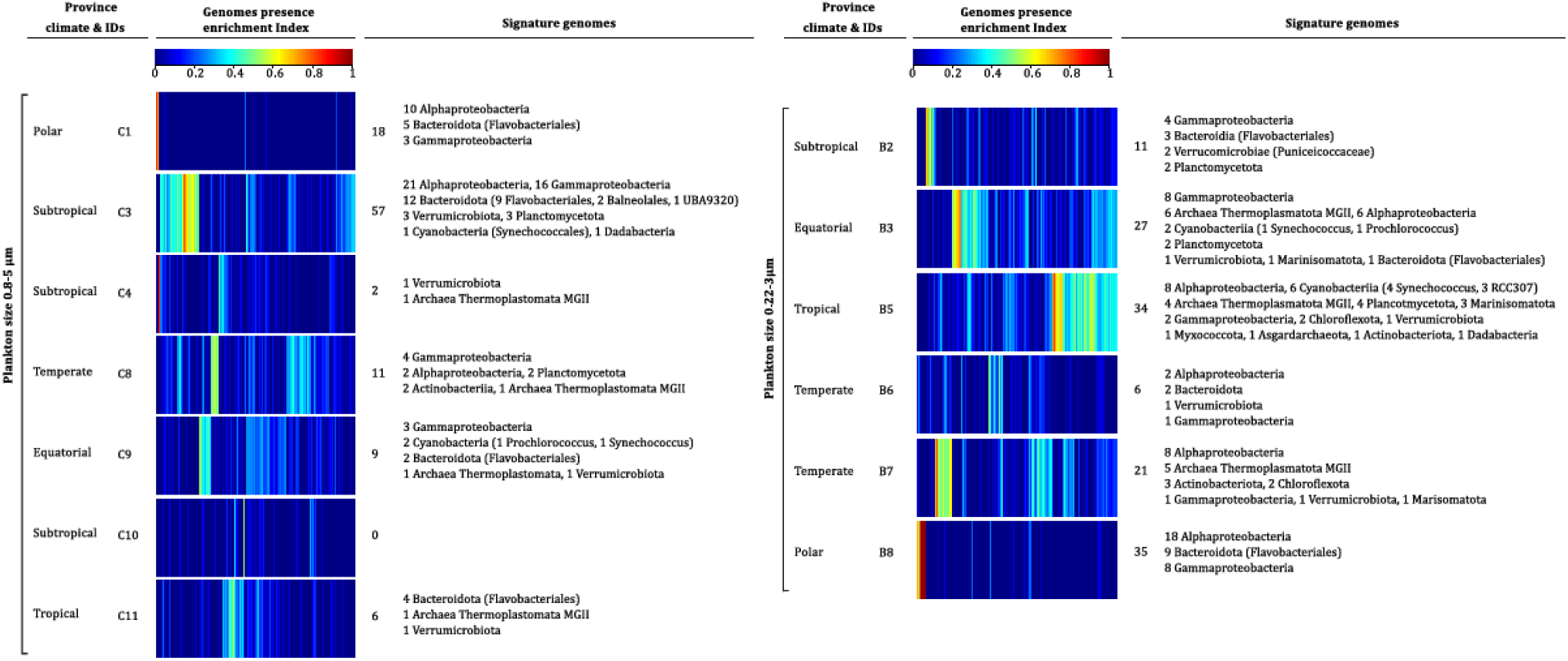
Prokaryotic signature genomes of provinces of the prokaryote (0.22-3 μm) and protist (0.8-5 μm) enriched size classes. Indexes of presence enrichment^51^ for 1888 genomes of prokaryotic plankton^32^ in corresponding provinces are clustered and represented in a color scale. Signature genomes (see *Methods*) are found for almost all provinces, their number and taxonomies are summarized (detailed list in Supplementary Table 6).

**Extended Data 3.**
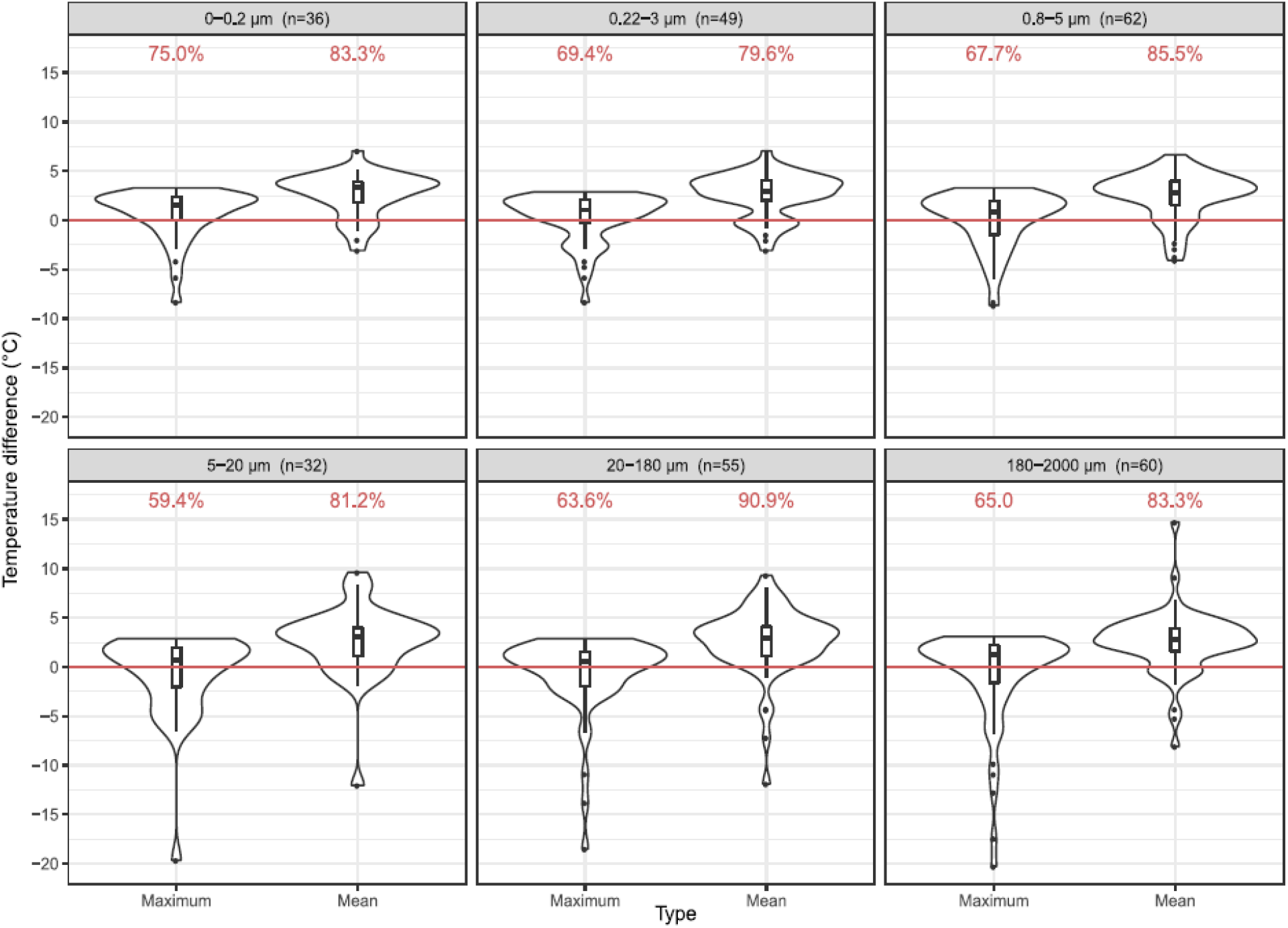
Distribution of deltas between future temperature at each sampling site (surface) minus either the mean or maximum temperature within their contemporary genomic province. For most of the sites and across size fractions the future temperature projected by the bias adjusted ESM ensemble model is higher than both the maximum and mean contemporary temperature of their genomic province.

**Extended Data 4.**
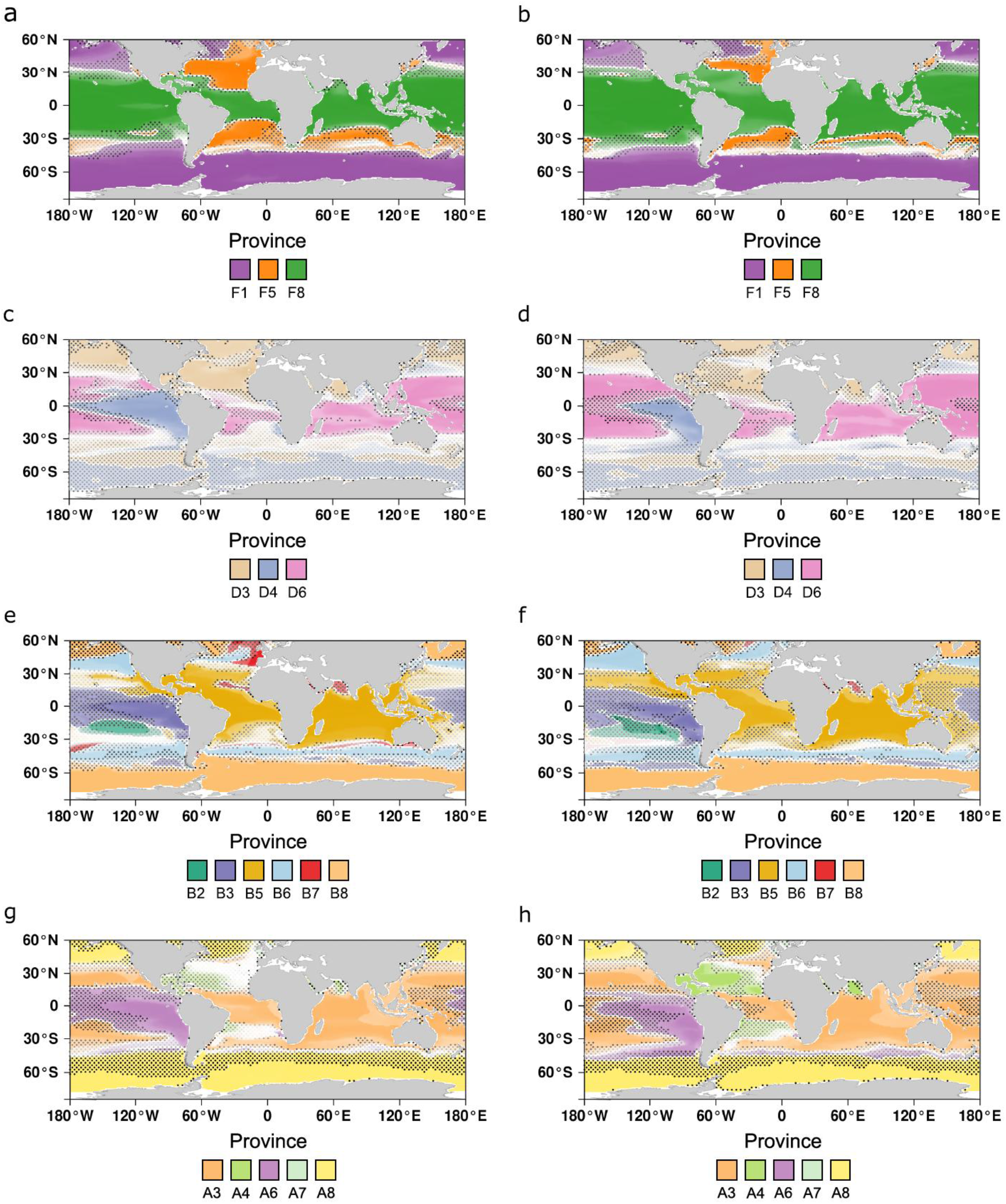
Global geographical patterns for 180-2000, 5-20, 0.22-3, 0-0.2 μm plankton size fractions in present day (a, c, e, g) and at the end of the century (b, d, f, h). The dominant province *i.e.* the one predicted to have the highest probability of presence is represented at each grid point of the map. The color transparency is the probability of presence of the dominant province. Expansion of tropical provinces and shrinkage of temperate communities is consistently projected in all size fractions.

**Extended Data 5.**
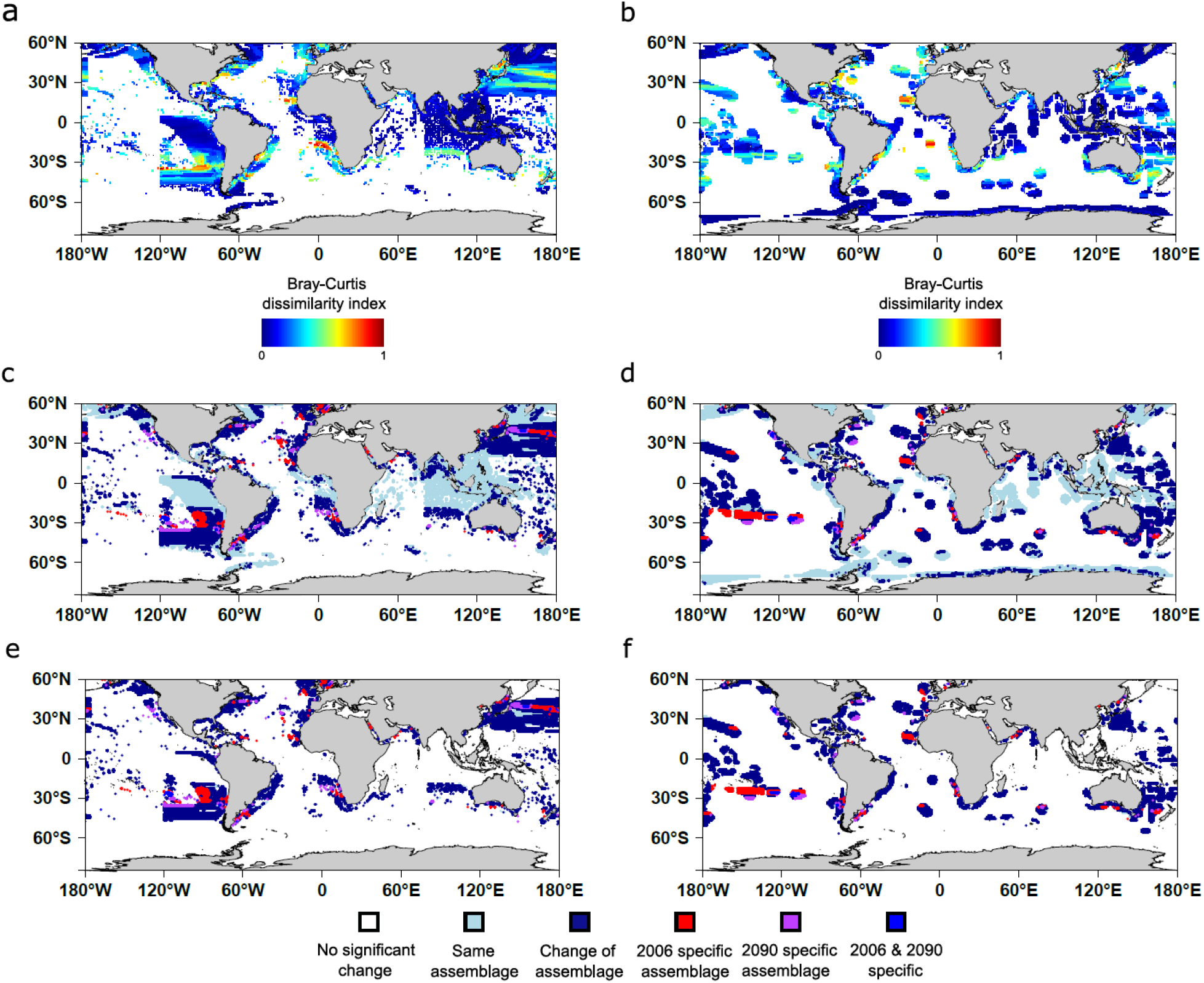
Bray-Curtis dissimilarity index and assemblage change maps comparing present day with end of the century projections of *dominant* provinces in (a, c, e) principal fisheries (4 last deciles^71^) and (b, d, f) Exclusive Economic Zones^72^. Assemblage changes in (**c**) Principal fisheries (**d**) Exclusive Economic Zones. Assemblage changes in (**e**) Principal fisheries (**f**) Exclusive Economic Zones with a Bray-Curtis dissimilarity index superior to 1/6.

**Extended Data 6.**
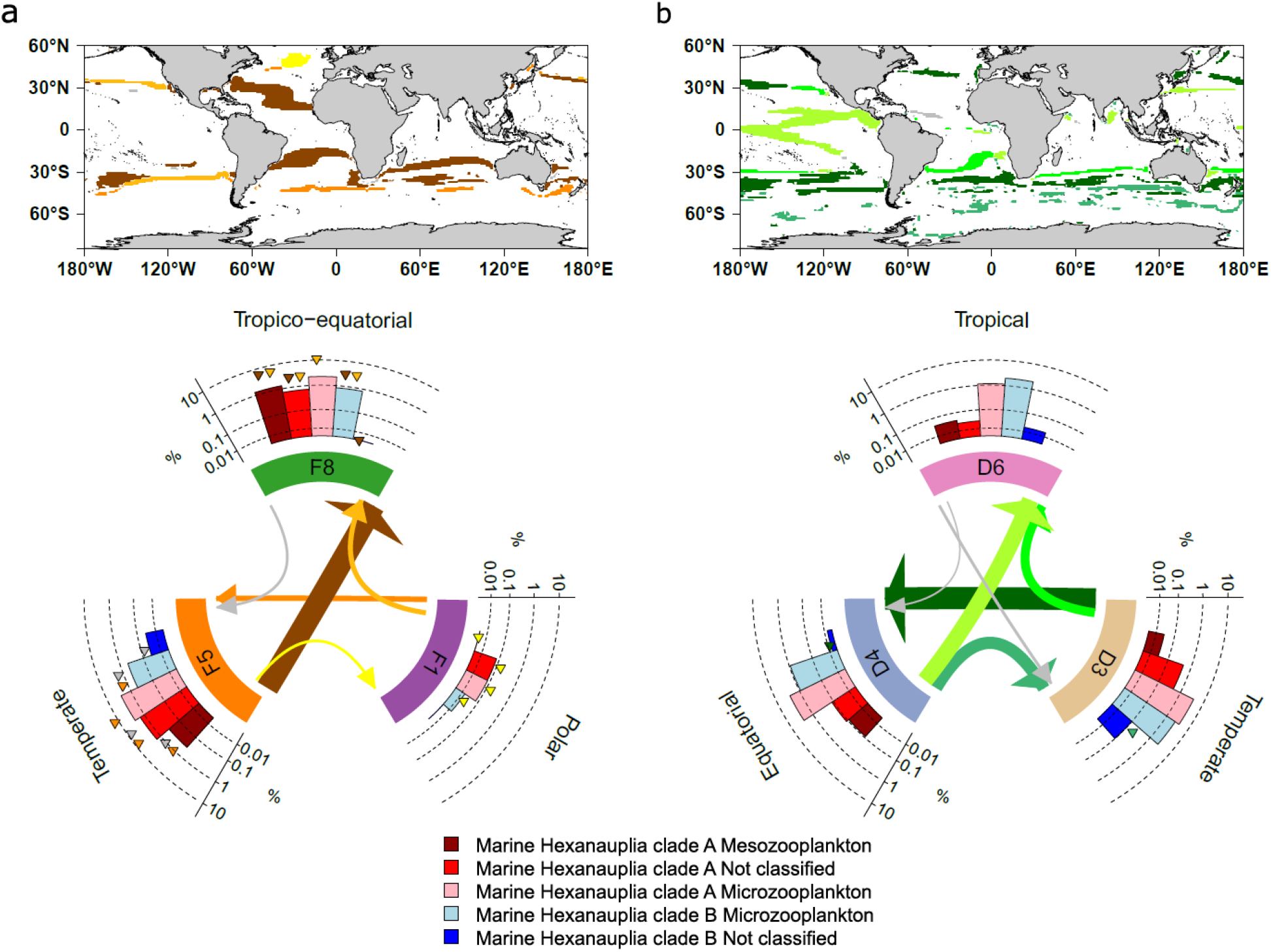
Projected compositional changes in marine hexanauplia in areas of dominant community change in size fraction (a) 180-2000 μm and (b) 5-20 μm. Top: Locations of dominant community change are highlighted using different colors depending on the dominant community transition. Bottom: Circular plots summarizing significant compositional shifts in marine hexanauplia: each type of transition is colored differently and according to the one on the map or in grey if they represent less than 2% of the transitions. Barplots represent mean relative abundances. Arrows represent dominant community shifts pointing towards the end of the century projected province and their widths are proportional to the area of change. Significant compositional changes in a type of genome are represented by triangles of the associated transition color.

**Extended Data 7.**
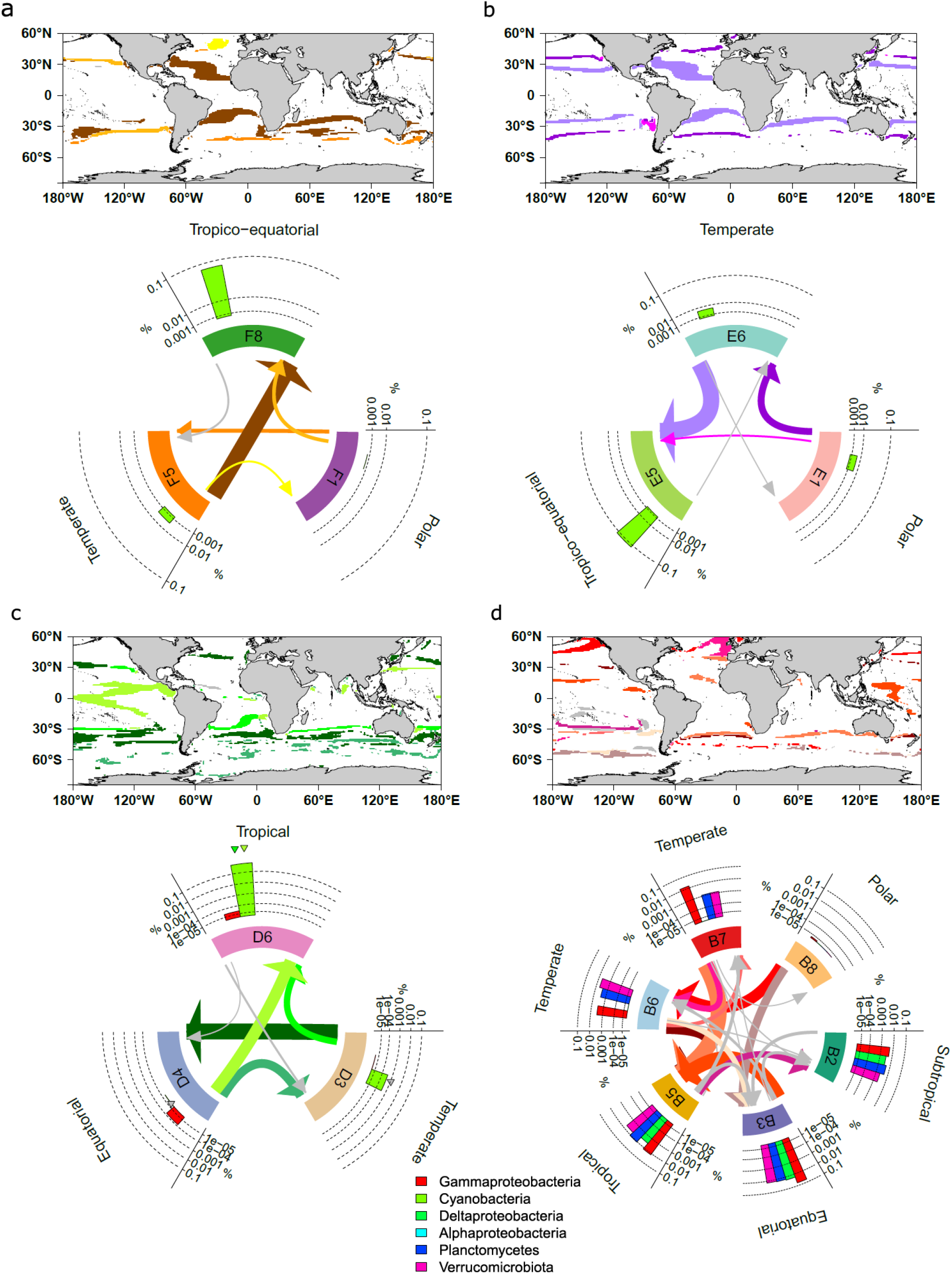
Projected compositional changes in bacterial diazotrophs in areas of dominant community change in size fraction (a) 180-2000 μm (b) 20-180 μm (c) 5-20 μm and (d) 0.22-3 μm. Top: Locations of dominant community change are highlighted using different colors depending on the dominant community transition. Bottom: Circular plots summarizing significant compositional shifts in marine diazotrophs: each type of transition is colored differently and according to the one on the map or in grey if they represent less than 2% of the transitions. Barplots represent mean relative abundances. Arrows represent dominant community shifts pointing towards the end of the century projected province and their widths are proportional to the area of change. Significant compositional changes in a type of genome are represented by triangles of the associated transition.

**Extended Data 8.**
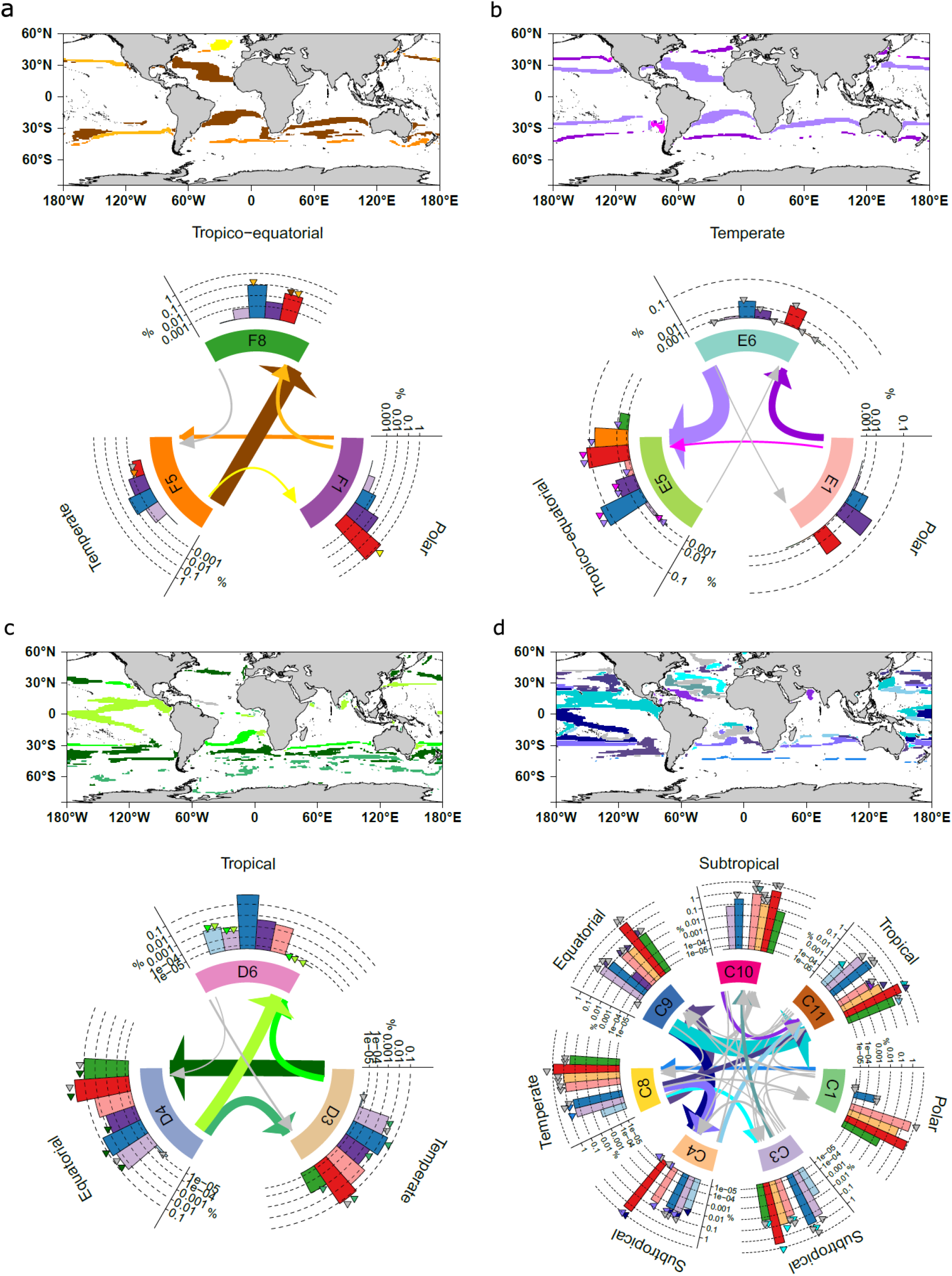
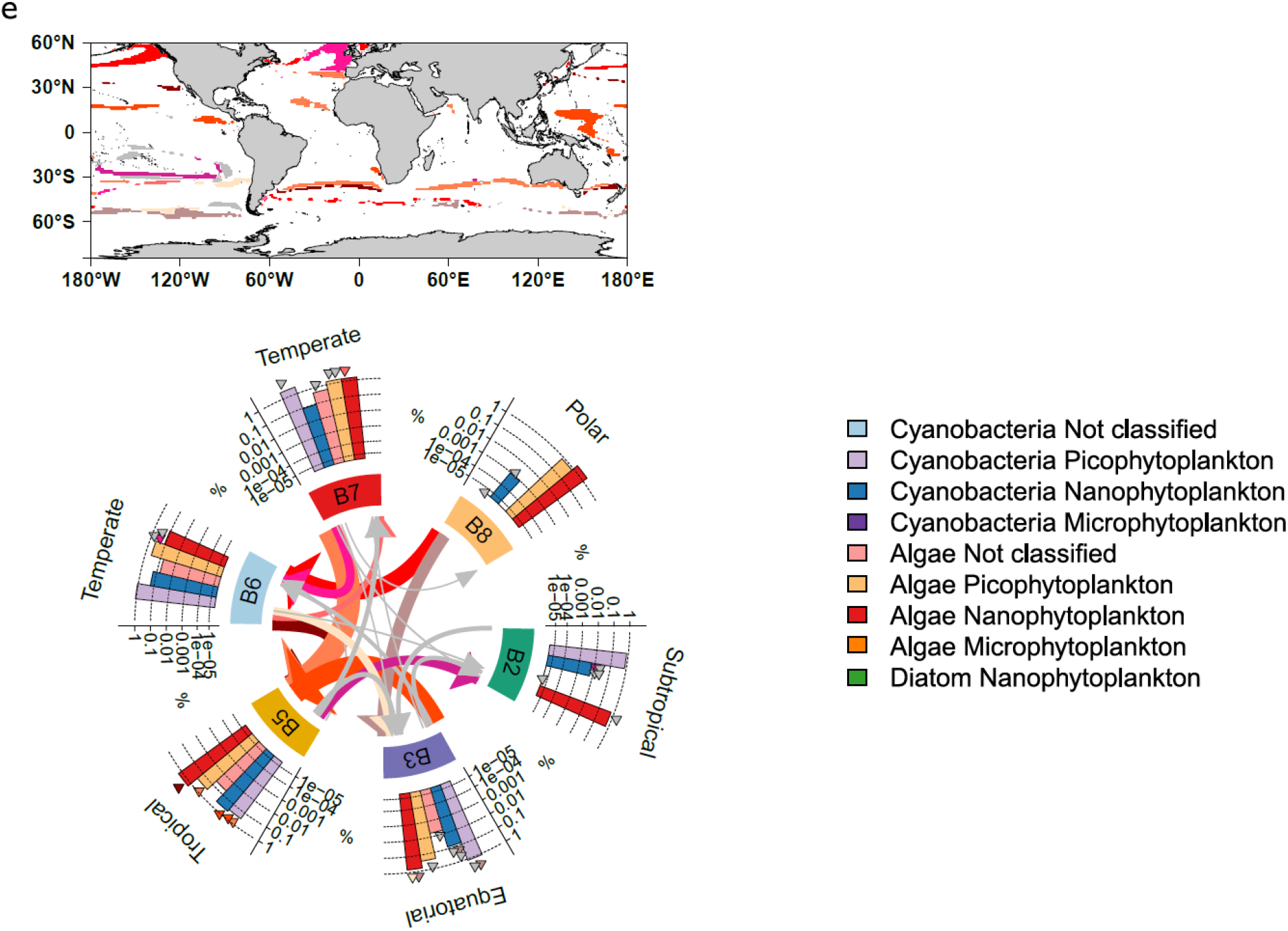
Projected compositional changes in phototrophs in areas of dominant community change in size fraction (a) 180-2000 μm (b) 20-180 μm (c) 5-20 μm (d) 0.8-5 μm and (e) 0.22-3 μm. Top: Locations of dominant community change are highlighted using different colors depending on the dominant community transition. Bottom: Circular plots summarizing significant compositional shifts in marine phototrophs: each type of transition is colored differently and according to the one on the map or in grey if they represent less than 2% of the transitions. Barplots represent mean relative abundances. Arrows represent dominant community shifts pointing towards the end of the century projected province and their widths are proportional to the area of change. Significant compositional changes in a type of genome are represented by triangles of the associated transition.

**Extended Data 9.**
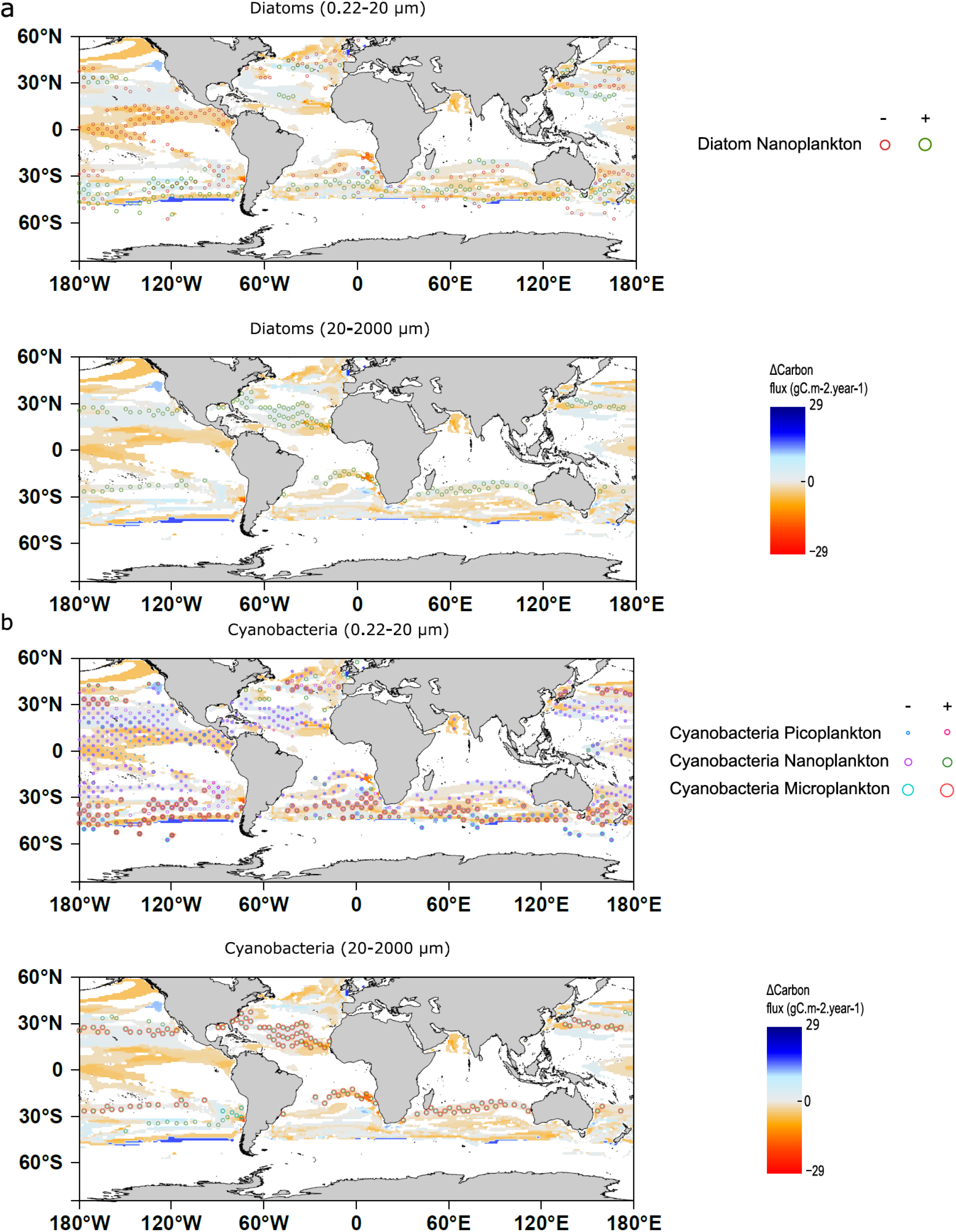
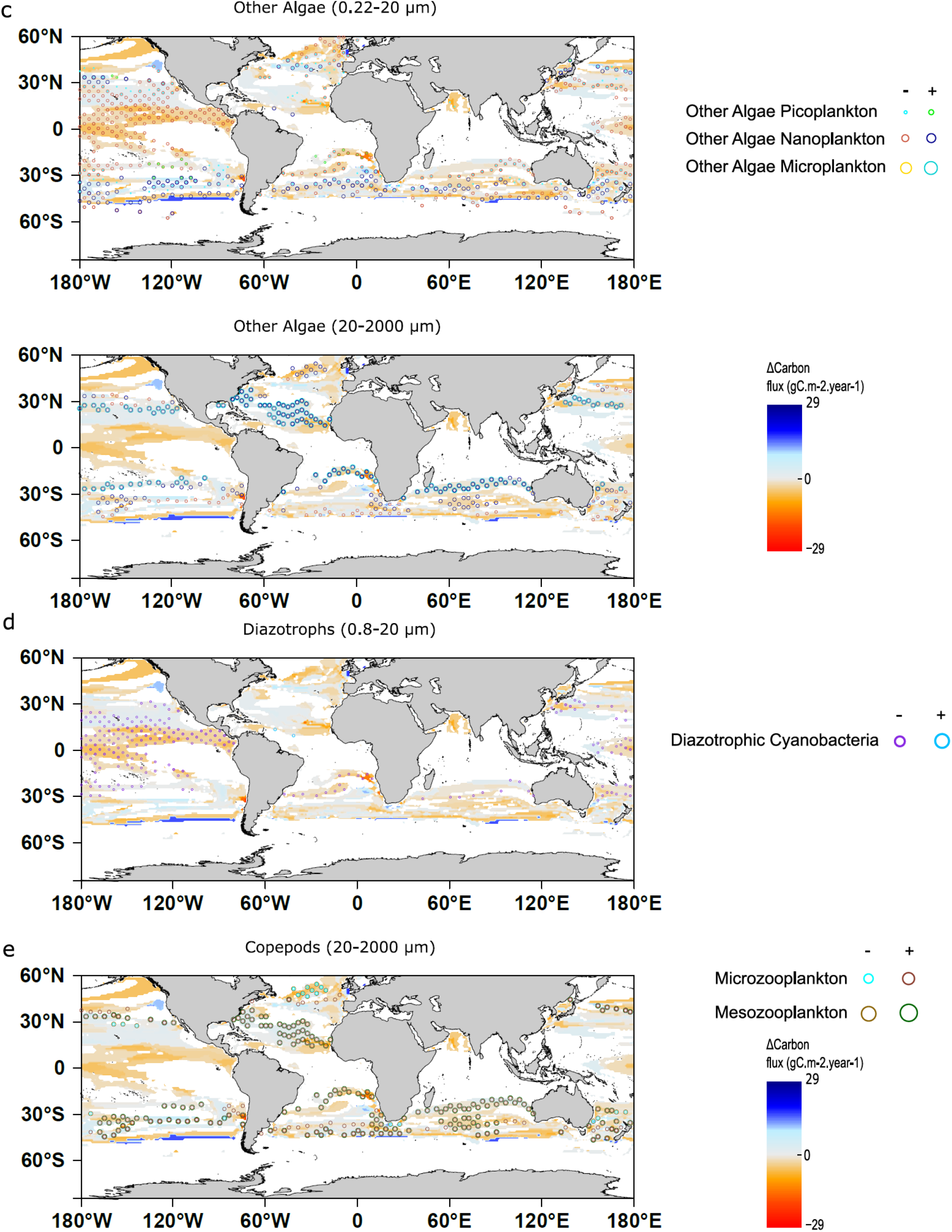
Maps of carbon export flux changes in link with organisms’ projected changes. Significant composition changes based on genomes relative abundances are represented for phototrophs, marine nitrogen fixers (Diazotrophic cyanobacteria) and copepods. For each map, transitions from several characteristic size classes are represented (**a**) Top: Diatoms 0.22-20 μm. Bottom: Diatoms 20-2000 μm (**b**) Top: Cyanobacteria 0.22-20 μm. Bottom: Cyanobacteria 20-2000 μm (**c**) Top: Other Algae 0.22-20 μm. Bottom: Other Algae 20-2000 μm (**d**) Diazotrophs 0.8-20 μm (**e**) Copepods 20-2000 μm.

**Extended Data 10.**
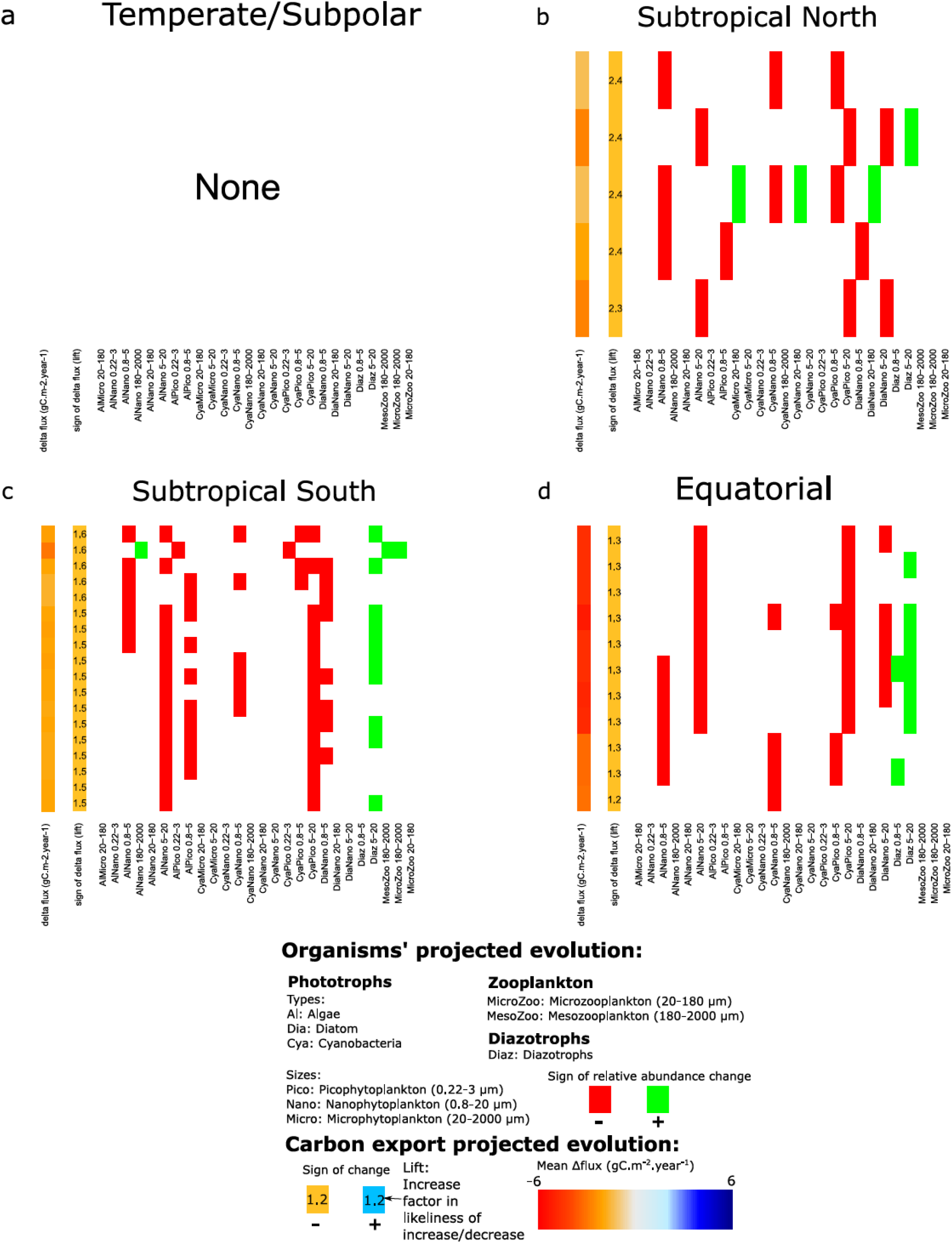
Association maps between carbon flux and changes in organism relative abundances. Each line represents an association rule between a change in carbon export found by the Apriori algorithm^45^ (first column: mean change in carbon export and second column: sign of the change and lift of the rule). The other columns represent the changes in community composition (red: decrease of the given group, green:increase) associated with this change in carbon export.

**Supplementary information 1 Niche models of genomic provinces**

To compute and test the validity of realized environmental niches, we train four machine learning techniques to probabilistically associate genomic provinces with environmental data: sea surface temperature, salinity, three macronutrients (dissolved silica, nitrate and phosphate), one micronutrient (dissolved iron) plus a seasonality index of nitrate. The four machine learning techniques are Gradient Boosting Machine (gbm)^1^, Random Forest (rf)^2^, single hidden layer Neural Networks (nn)^3^ and Generalized Additive Models (gam)^4^. Following a cross-validation framework, a valid environmental niche is obtained for 27 out of 38 initial provinces (71%) comforting their definition and covering 529 samples out of 595 (89%, Supplementary Fig. 1). Rejected provinces contain relatively few stations (mean of 6 ± 2.6 versus 19 ± 15.3 for valid provinces, p-value<10^-3^ Wilcoxon test^5^). For spatial and temporal extrapolations of the provinces presented below, we use the ensemble model approach^6^ that considers mean predictions of machine learning techniques.

**Supplementary information 2 Integrated biogeographies using the PHATE algorithm**

We use the PHATE dimension reduction algorithm^7^ to combine all provinces for all size classes into a single consensus biogeography revealing 4 or 7 robust clusters (Supplementary Fig. 3, *methods*). The 4 cluster consensus biogeography is mainly latitudinally organized distinguishing polar, subpolar, temperate and tropico-equatorial regions. The 7 cluster consensus biogeography distinguishes the equatorial pacific upwelling biome and three subpolar biomes that most likely reflect the chemico-physical structuring of the Southern Ocean and known polar fronts^8^ (red lines Fig. 2h). However, learning data are scarcer south of 60°S so these extrapolations need to be taken with caution.

PHATE is also used for modeled projections into two comparable consensus maps, one for present day and one for the end of the century (Supplementary Fig. 13). Some particularly visible patterns of geographical reorganization are common to several or even all size fractions and visible when comparing the two consensus maps (Supplementary Fig. 13 compared to Fig. 3a-d and Extended Data 4). For example, the tropico-equatorial and tropical provinces expand in all size fractions and the provinces including the pacific equatorial upwelling shrink for size fractions smaller than 20 µm.

**Supplementary information 3 Comparison of the biogeographies with existing partitions of the oceans**

Previous ocean partitioning either in biomes^9–11^ or biogeochemical provinces (BGCPs)^10^ are based on physico-biogeochemical characteristics including SST^9–11^, chlorophyll *a*^9–, 11^, salinity^9–11^, bathymetry^9, 10^, mixed layer depth^11^ or ice fraction^11^. Considering three of these partitions as examples we notice differences with our partitions (Supplementary Fig. 6-7) for example in terms of the number of regions in the considered oceans (56 for 2013 Reygondeau et al. BGCPs^10^, 17 for Fay and McKingley^11^) and their structure (the coastal biome for 2013 Reygondeau et al. biomes^10^). Numerical comparison of our partitions with others (*Methods*) reveals low similarity between them, the highest being with Reygondeau biomes (Supplementary Figs. 6-8). However, biomes and BGCP frontiers closely match our province frontiers in many cases. Near the frontiers, *dominant* provinces have smaller probabilities in agreement with smooth transitions instead of sharp boundaries as already proposed^9^ and with a seasonal variability of the frontiers^10^ (Supplementary Fig. 6). Some of these transitions are very large and match entire BGCPs, for example in subtropical North Atlantic and subpolar areas where high annual variations are well known^10^.

**Supplementary information 4 Shifts of the *climato-genomic* provinces in response to climate change**

Centroids of provinces with *dominance* areas larger than 10^6^ km^2^ within a basin would be moved at least 200 km away for 77% of them, 96% of which move poleward (Supplementary Figs. 11 and 12). While a few longitudinal shifts larger than 1000 km are projected, the distribution of latitudinal shifts is largely concentrated around the mean (290 km) with no shifts superior to 1000 km (Supplementary Fig. 12b). These important longitudinal shifts corroborate existing projections^12–14^ and differ from trivial poleward shifts due to temperature increase, reflecting more complex spatial rearrangements of the other environmental drivers (Supplementary Fig. 10). The average displacement speed of the provinces’ centroids is 76 ± 79 km.dec^-1^ (latitudinally mean of 34 ± 82 km.dec^-1^, longitudinally 59 ± 82 km.dec^-1^).

Projected shifts in phytoplankton enriched provinces corroborate previously published shifts of North Pacific phytoplankton biomes: provinces C4 and C9 are projected to shift respectively at speeds of 118 km.dec^-1^ and 195 km.dec^-1^ comparable to 100 km.dec^-1^ and 200 km.dec^-1^ for the subtropical and tropical biomes of Polovina et al.^14^.

For all size fractions, climate change would lead to a poleward expansion of tropical and equatorial provinces at the expense of temperate provinces (Supplementary information 4, Supplementary Table 2, Extended Data 7, Supplementary Fig. 13). This is illustrated by the temperate province F5 of size fraction 180-2000 (Supplementary Fig. 12), which is projected to shrink in the five major basins. In the North Atlantic, its centroid would move approximately 800 km to the northeast (Supplementary Fig. 12c). Similar trends are found comparing present day and end of the century consensus maps (Supplementary information 2, Supplementary Fig. 13).

**Supplementary information 5 Expansion and shrinkage of provinces in response to climate change**

To quantify patterns of expansion or shrinkage of the provinces, we calculate the surface covered by the *dominant* provinces weighted by probabilities of presence (Supplementary Fig. 14, Supplementary Table 2). In this way, *dominant* provinces are defined on 100% of the surface ocean (327 million km^2^) but their presence probabilities correspond to the equivalent of 45 to 74% (due to sampling variability and niche overlaps) of the surface ocean depending on the plankton size fraction (Table 1). Overall, our results indicate expansions of the surface of tropical and tropico-equatorial provinces but in very different ways depending on the size fractions of organisms. The surface area of temperate provinces is ∼22 million km^2^ on average (from 10 Mkm^2^ for 0-0.2 µm to 49 Mkm^2^ for 20-180 µm) and should decrease by 36% on average (from - 20 % for 5-20 µm up to -54% for 0.8-5 µm, -12 million km^2^ on average, +6% for 0.22-3 µm). Tropical provinces cover ∼118 million km^2^ on average (from 86 Mkm^2^ for 0.8-5 µm up to 169 Mkm^2^ for 180-2000 µm) and their coverage should increase by 32% on average (from +13% for 0-0.2 µm up to +75% for 0.8-5 µm, +25 million km^2^ on average) (Supplementary Fig. 14 and Supplementary Table 2).

**Supplementary information 6 Changes in the distribution of grazers and phototrophs**

Copepods are cosmopolitan small crustaceans. These abundant grazers contribute to the biological pump and their body size is considered to be a key trait for carbon export^15^. They feed on smaller plankton^15^ and are prey to higher trophic levels. We characterize the composition of provinces using 198 environmental genomes detected in at least 5 samples and annotated as Marine Hexanauplia of clade A and B. They are further divided into five subgroups based on their differential abundances across the large size classes (>5 μm): 8 mesozooplankton clade A, 19 unclassified clade A, 71 microzooplankton clade A, 10 unclassified clade B and 90 microzooplankton clade B. Therefore, large copepods are preferentially from clade A in our data.

The relative abundances of unclassified clade B in size class 20-180 μm (Fig. 3e) and of mesozooplankton clade A in size class 180-2000 μm, (Extended Data 6a) are projected to increase in regions where the temperate province is replaced by the tropico-equatorial province. In areas where the polar province is replaced by the temperate province, a greater relative abundance in microzooplankton clade A (1% to 10%) (Fig. 3e) is projected. No significant compositional differences in the provinces of size fraction 5-20 μm are found (Supplementary Fig. 6b). These results highlight potential significant compositional shifts by the end of the century in the different clades and sizes of main grazers.

Phototrophs are the primary producer of the ocean that fix carbon from the atmosphere via photosynthesis and therefore sustain the whole trophic chain and carbon export to the deep oceans^16–18^. Their size is a key trait for carbon export: picophytoplankton (between 0.2 and 2 µm) and nanophytoplankton (2-20 µm) are believed to contribute relatively little to carbon export^19^, although the question is under debate^20^, while larger cells of microphytoplankton (>20 µm) especially diatoms are large contributors^21^. We characterize our genomic provinces by their content in 231 phototrophs annotated by their preferential size (nano to microphytoplankton) and taxonomy (diatoms, cyanobacteria and other algae (referred to as ‘algae’)).

In regions of temperate to tropico-equatorial transition, nano-algae are projected to increase in size fraction 180-2000 μm (Extended Fig. 8a) while micro-algae, nano-diatoms and cyanobacteria (both nano- and micro-) are projected to increase in size fraction 20-180 µm (Extended Fig. 8b). In size fraction 5-20 μm, nano-diatoms, algae and cyanobacteria are all projected to increase in temperate to equatorial transitions while nano-diatoms and algae and pico-cyanobacteria decrease in equatorial to tropical transitions (Extended Fig. 8c). In size fraction 0.8-5 μm, nano-algae are decreasing in equatorial to tropical transitions. Nano-diatoms and cyanobacteria are decreasing in temperate to subtropical transitions (towards C4). Finally in size fraction 0.22-3 μm, in temperate to tropical transitions, pico-cyanobacteria and algae are projected to decrease. Overall, changes in phototroph composition are complex and our dataset is not sufficient to describe the entire diversity of phototrophs (especially large diatoms and dinoflagellates that can have bigger genomes).

---

**Supplementary Fig. 1.**
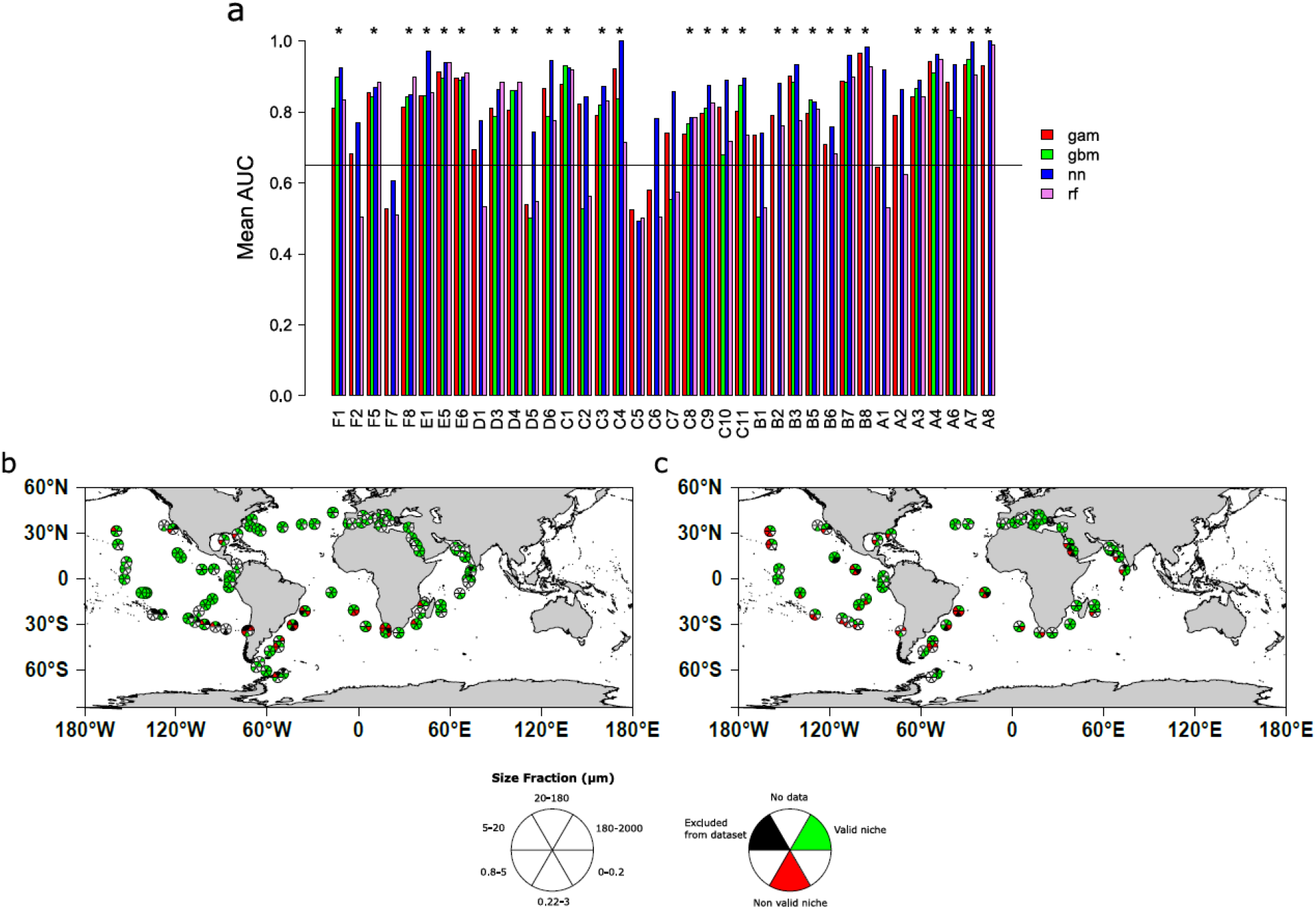
(a) Barplot of mean AUC over a random 30-fold cross validation process of the 38 initial metagenomic provinces. (b,c) Map of validated and non validated stations across the six size fractions in surface samples (b) and DCM samples (c). (**a**) Mean AUC (Area Under the receiver operating Characteristic)^22^ is plotted for the best hyperparameter combination of the 4 machine learning techniques used for each of the 38 metagenomic provinces. A star for valid models is drawn at the top of each considered valid model. (**b-c**) For each Tara sample present in the dataset at surface (**b**) or Deep Chlorophyll Maximum (DCM) (**c**) and for each size fraction, the filter (one size fraction at one location and one depth) belongs either to a validated niche (green), a non validated niche (red), an excluded filter (sea *Materials & Methods*) (black) or no data is available (white). Non validated niches represent only 11% of filters present in the dataset (66 out of 595).

**Supplementary Fig. 2.**
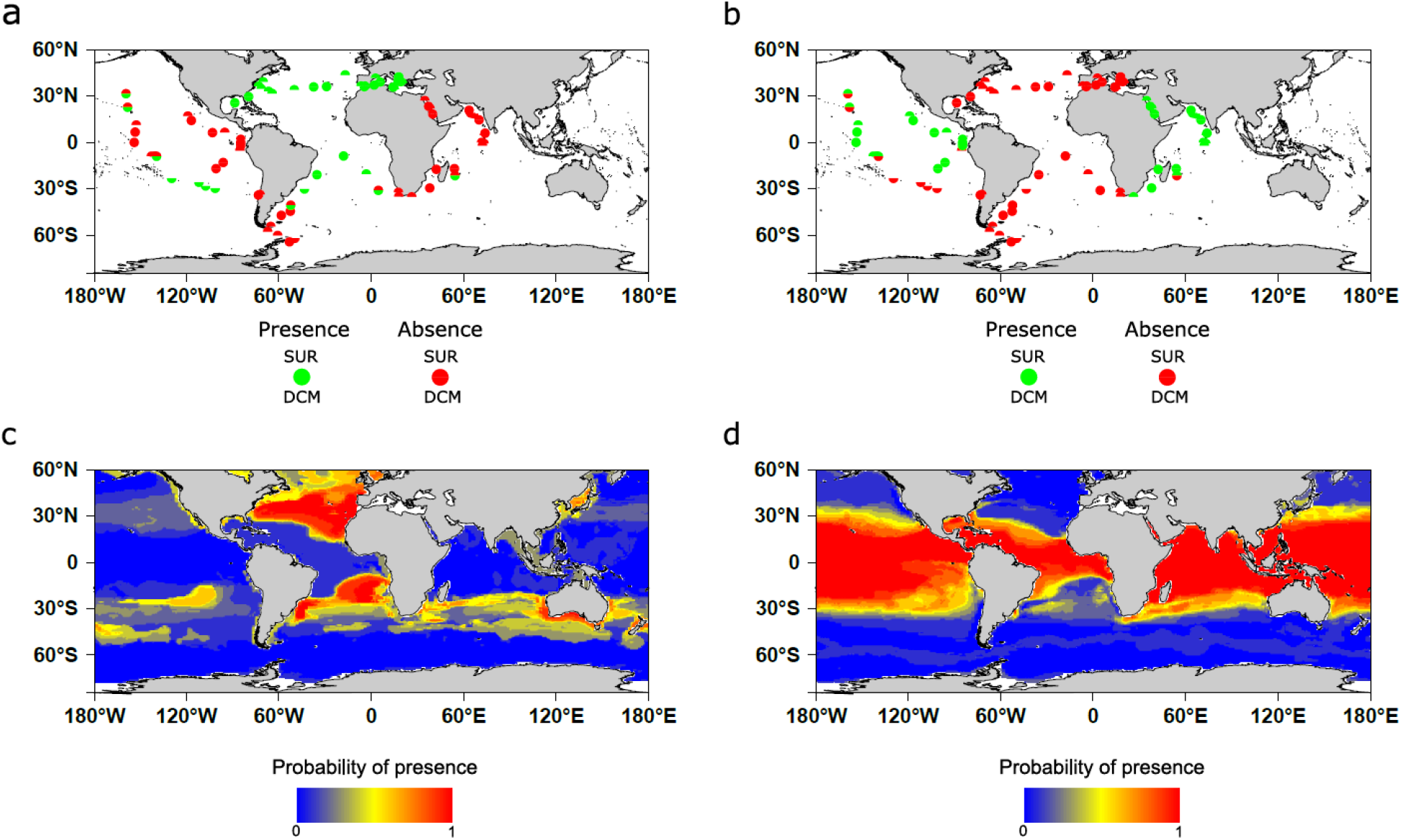
Sampling and projection maps on WOA13 climatological data of two example provinces from size fraction 180-2000 µm. (**a**) Sampling map of province F5. **(b)** Province F8. (**c**) Projection map of province F5 on WOA13^23^. (**d)** Province F8. Qualitatively, projection maps are coherent with sampling maps of the two provinces with the highest probability of presence projected in the sampling regions. Other presence zones are also predicted by the projection. Sampling of these zones might be interesting to confirm our approach and projections such as South of Australia where a high probability of presence for province F5 is predicted.

**Supplementary Fig. 3.**
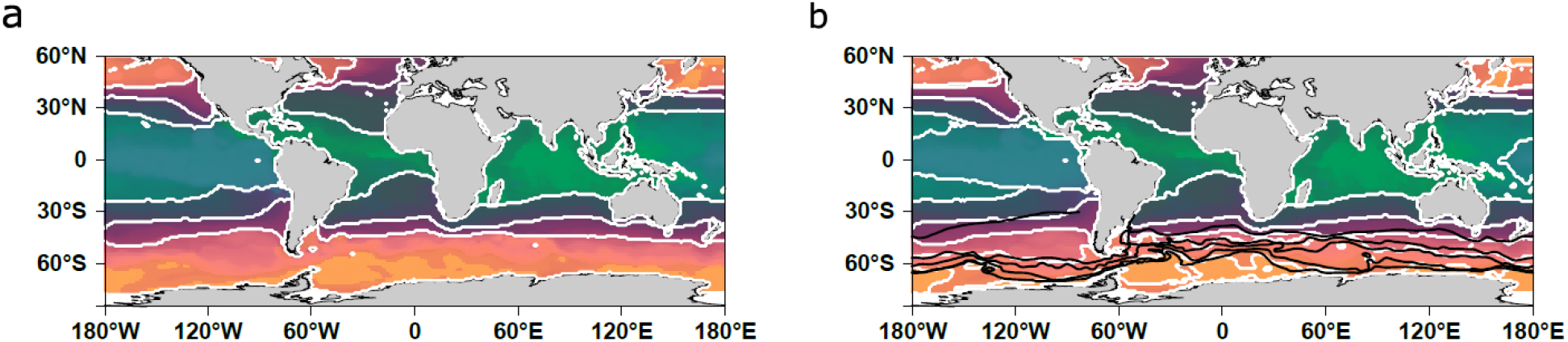
Present day combined size class biogeographies using the PHATE algorithm on WOA13 dataset. (**a**) Combined size class biogeography using PHATE algorithm^7^ partitioned in 4 clusters. Each grid point is associated to a color depending on its PHATE coordinates (using three axes). Each coordinate is respectively assigned to a given degree of color between 0 and 255 according to equation 2 (either red green or blue depending on the axis).(**b**) Combined size class biogeography using PHATE algorithm partitioned in 7 clusters overlaid with Antarctic Circumpolar Current boundaries (black). Colors are the same as in (**a**).

**Supplementary Fig. 4.**
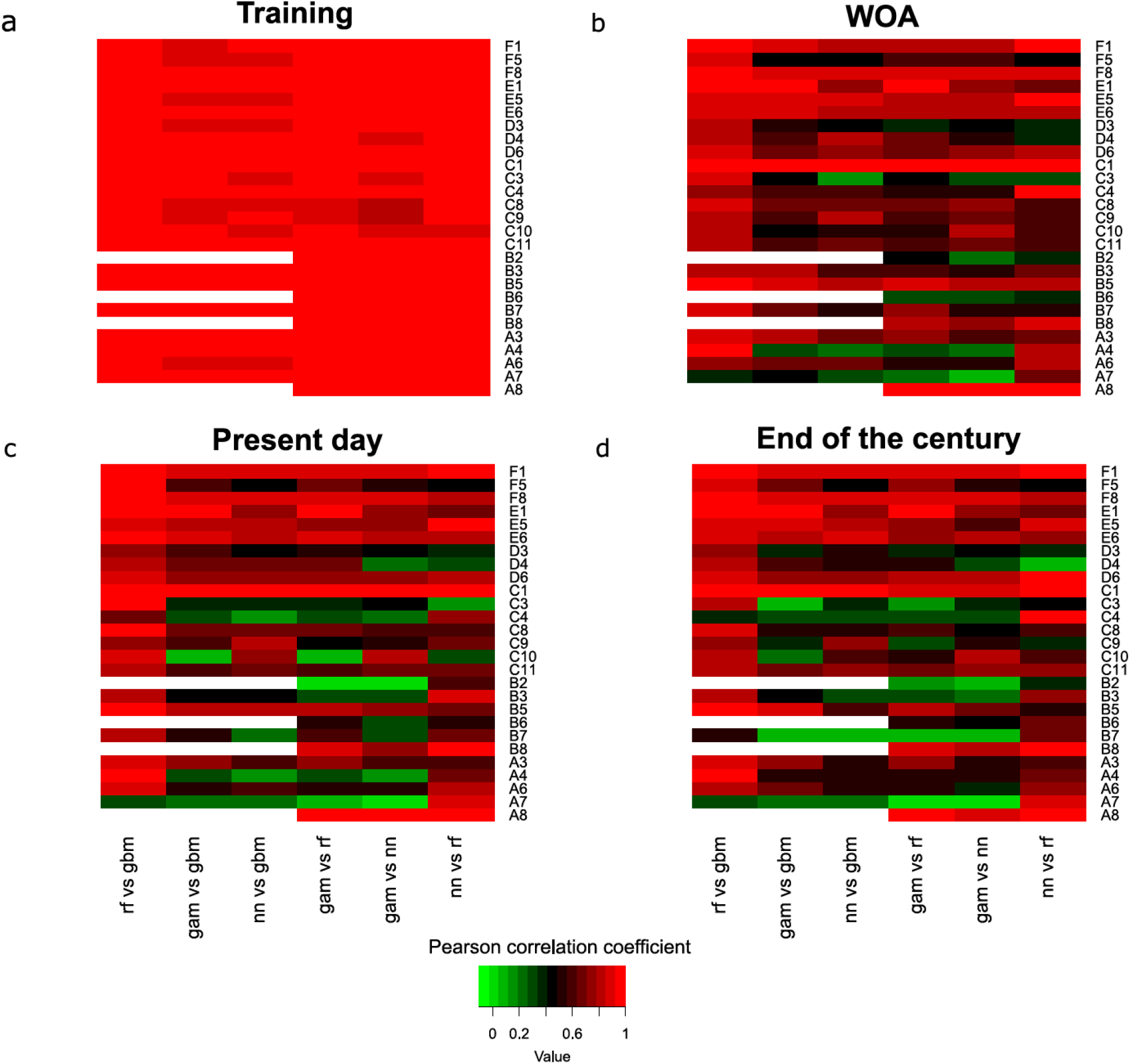
Pairwise Pearson correlation coefficient heat maps between outputs of the 4 machine learning models. rf: Random Forest, gbm: gradient boosting machines, gam: general additive models, nn: neural networks. The rows are the provinces of the different size fractions. The columns are the pairwise comparisons of each machine learning technique. (a) Training set outputs. (b) WOA13 average data projection outputs (except for Iron, PISCES-v2^24^ is used). (c) Present day data projection outputs (bias corrected mean model of 6 Earth System Models). (d) End of the century projection outputs. On the training set, models are in agreement with most of the correlation coefficients superior to 0.9. A drop in correlation is observed for modeled data (c, d) especially in small size fractions. This shows modeled data are more distant from the training set than WOA data. Random Forest and Gradient Boosting Machine are in very good agreement (first columns) which could be expected as they are both based on multiple decision trees.

**Supplementary Fig. 5.**
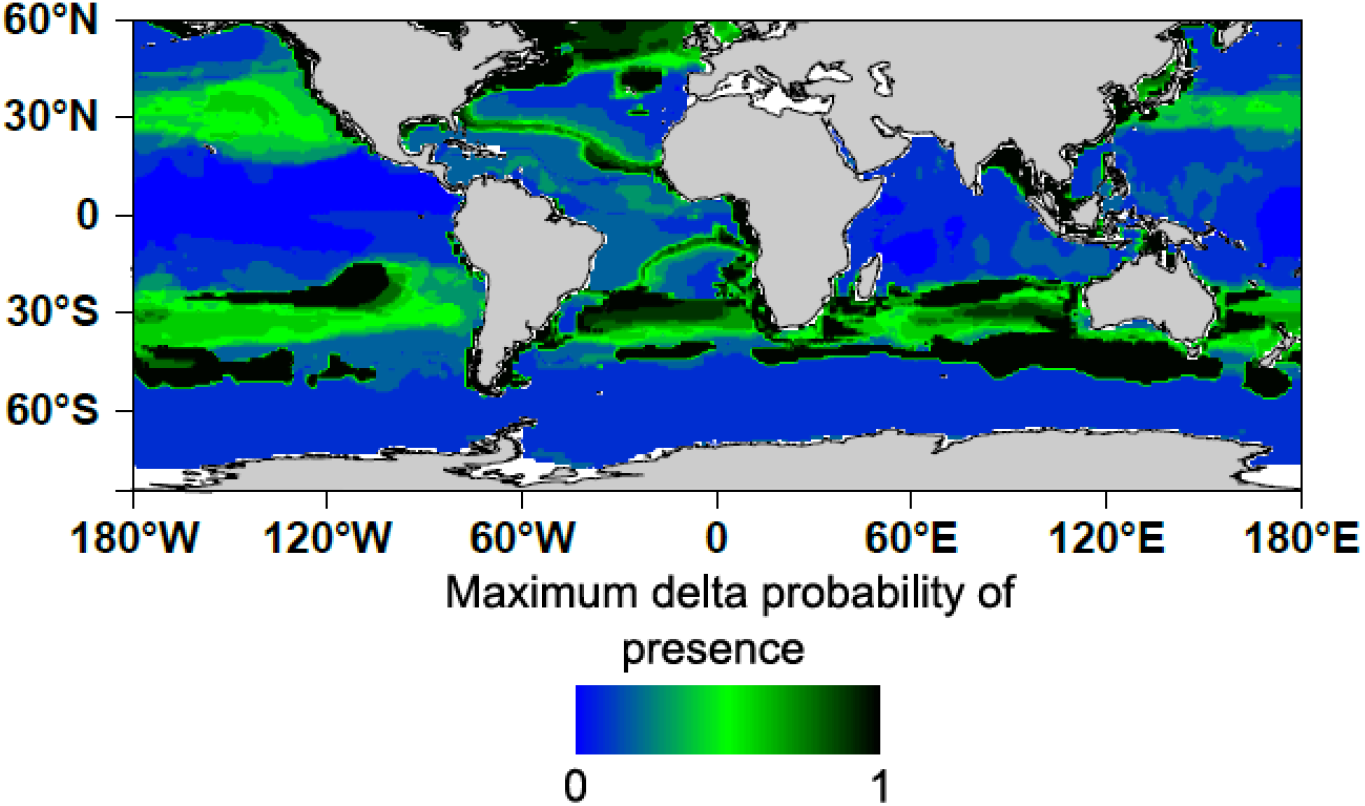
Probability range map (WOA data) of province F5 of size fraction 180-2000 µm. At each point of the grid, the maximum delta probability of presence between the 4 machine learning projections is calculated. In black are the zones where two models completely disagree: one model predicts presence with certainty whereas the other predicts absence with certainty. Disagreement appears mostly in uncertain presence areas (Supplementary Fig. 2c) whereas models are generally in good agreement in absence areas (blue zones).

**Supplementary Fig. 6.**
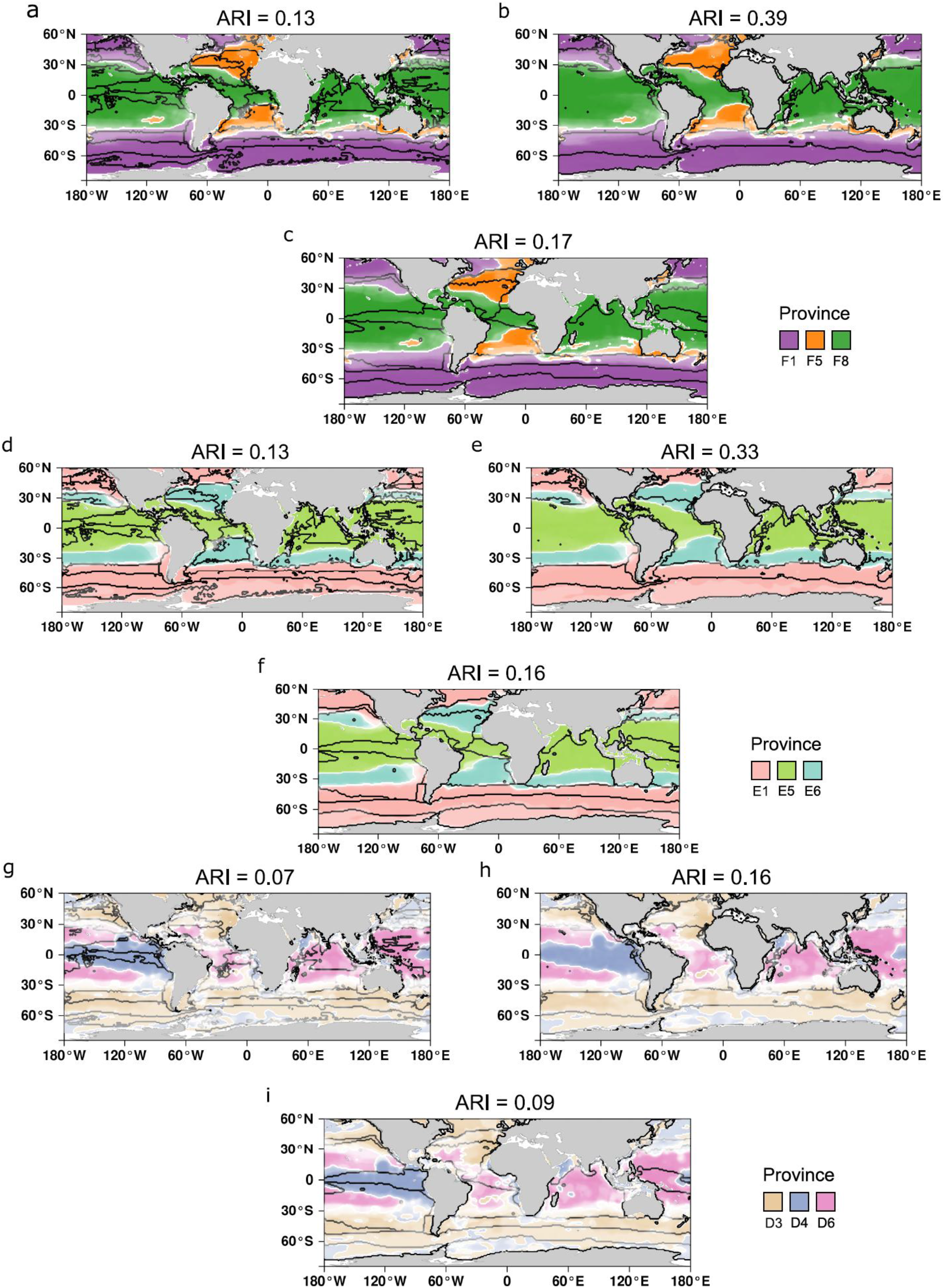
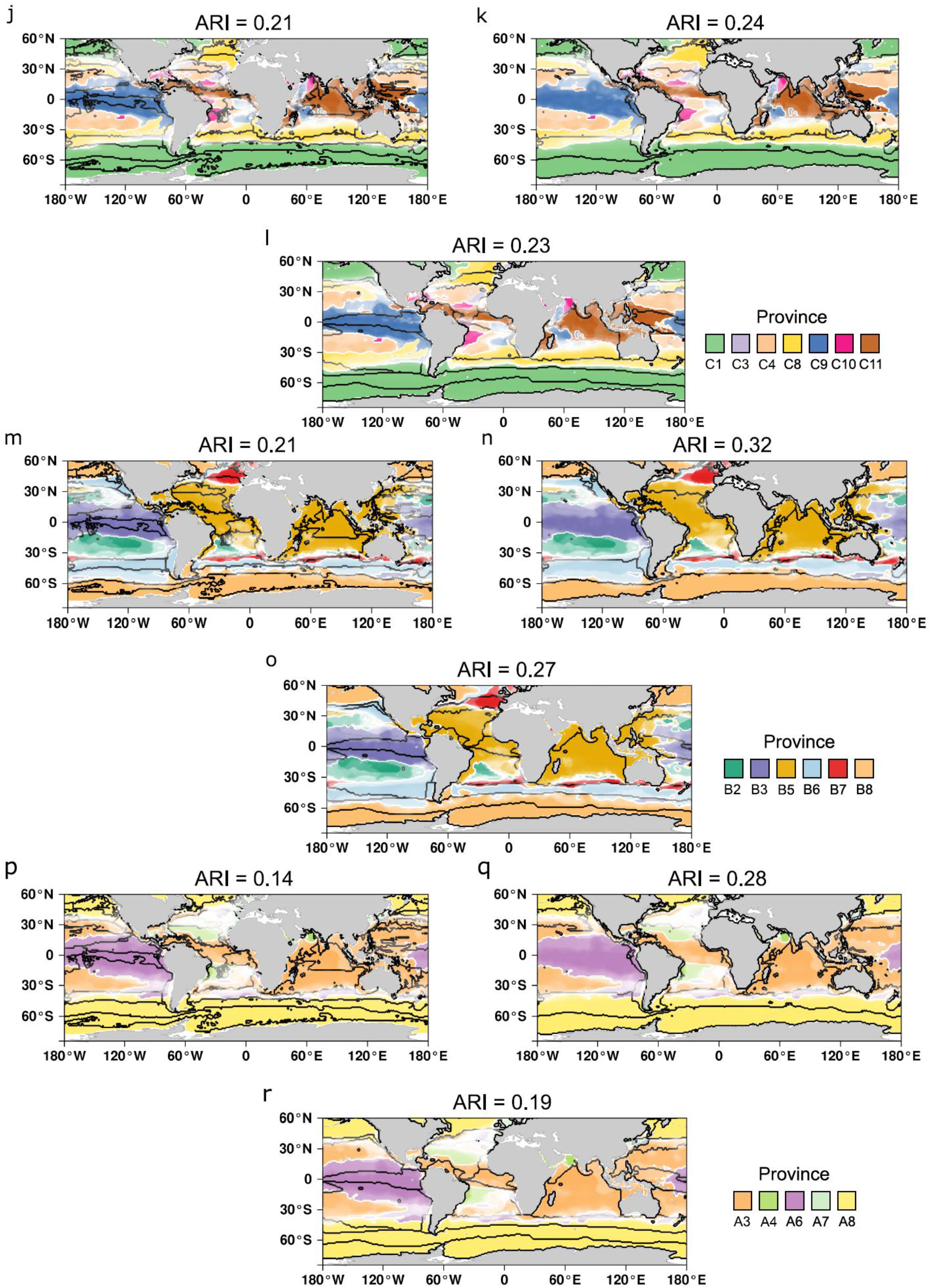
Three existing oceanic partitions overlaid on top of plankton provinces for the six fractions and combined size fractions. The three oceanic partitions are Reygondeau et al. Biogeochemical Provinces (BGCP)^10^; Reygondeau et al. Biomes^10^; Fay and McKingley Biomes^11^. Each partitioning mask is overlaid in the above order on top of plankton provinces for the six size fractions. (**a-c**) 180-2000 µm (**d-f**) 20-180 µm (**g-i**) 5-20 µm (**j-l**) 0.8-5 µm(**m-o**) 0.22-3 µm and (**p-r**) 0-0.2 µm. Above all maps, the adjusted rand index (ARI, an index used to compare partitions), for the comparision of mapped biogeographies with the black line mask is shown.

**Supplementary Fig. 7.**
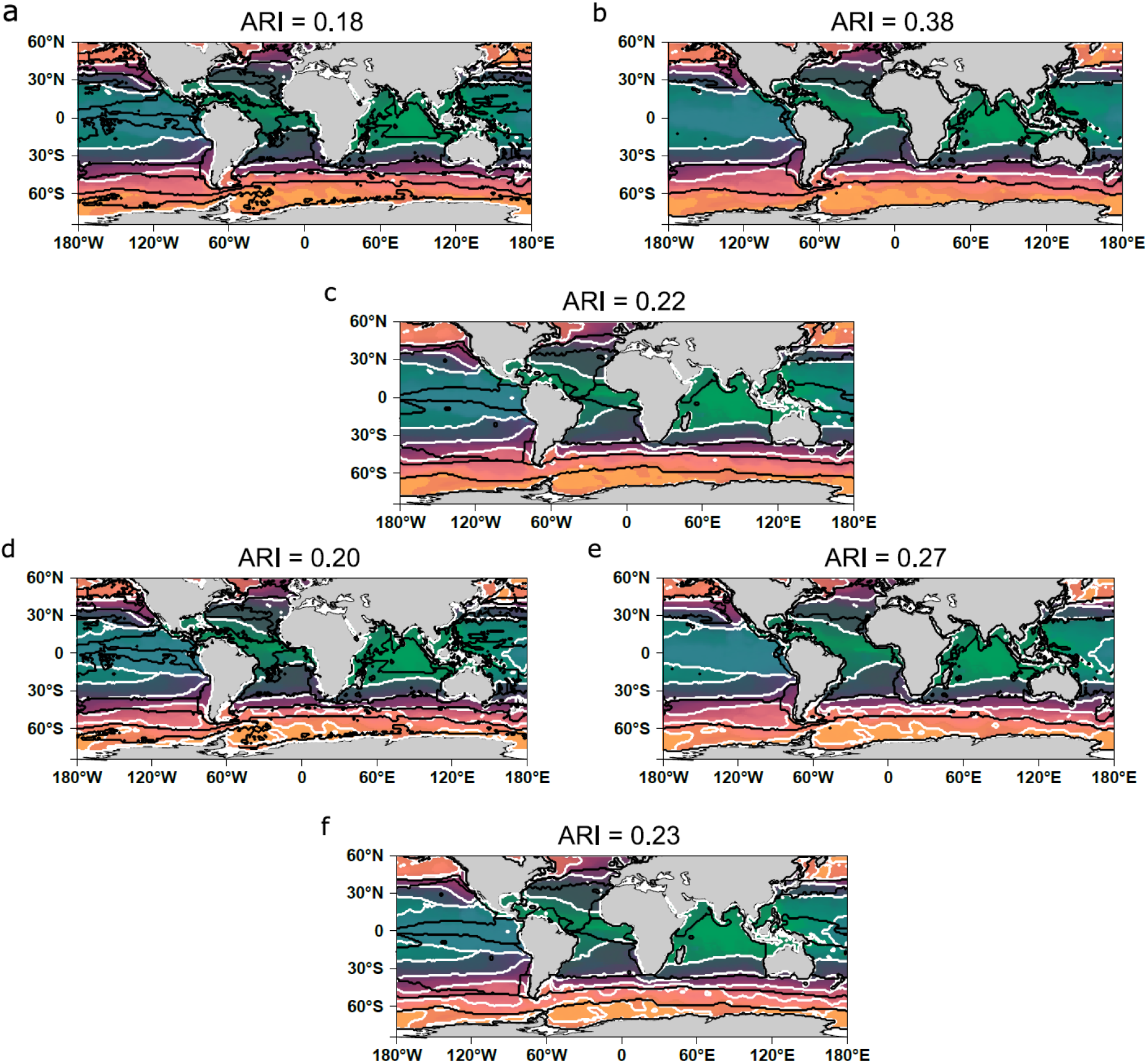
Three existing oceanic partitions overlaid on top of plankton provinces for the combined size fractions. The three oceanic partitions are Reygondeau et al.^10^ Biogeochemical Provinces (BGCP); Reygondeau et al. Biomes^10^; Fay and McKingley Biomes^11^. Each partitioning mask is overlaid in the above order on top of size fraction plankton provinces combined by the PHATE algorithm in (**a-c**) 4 clusters and (**d-f**) 7 clusters. Each grid point is associated with a color depending on its PHATE coordinates (using three axes). Each coordinate is respectively assigned to a given degree of color between 0 and 255 according to equation 2 (either red green or blue depending on the axis).

**Supplementary Fig. 8.**
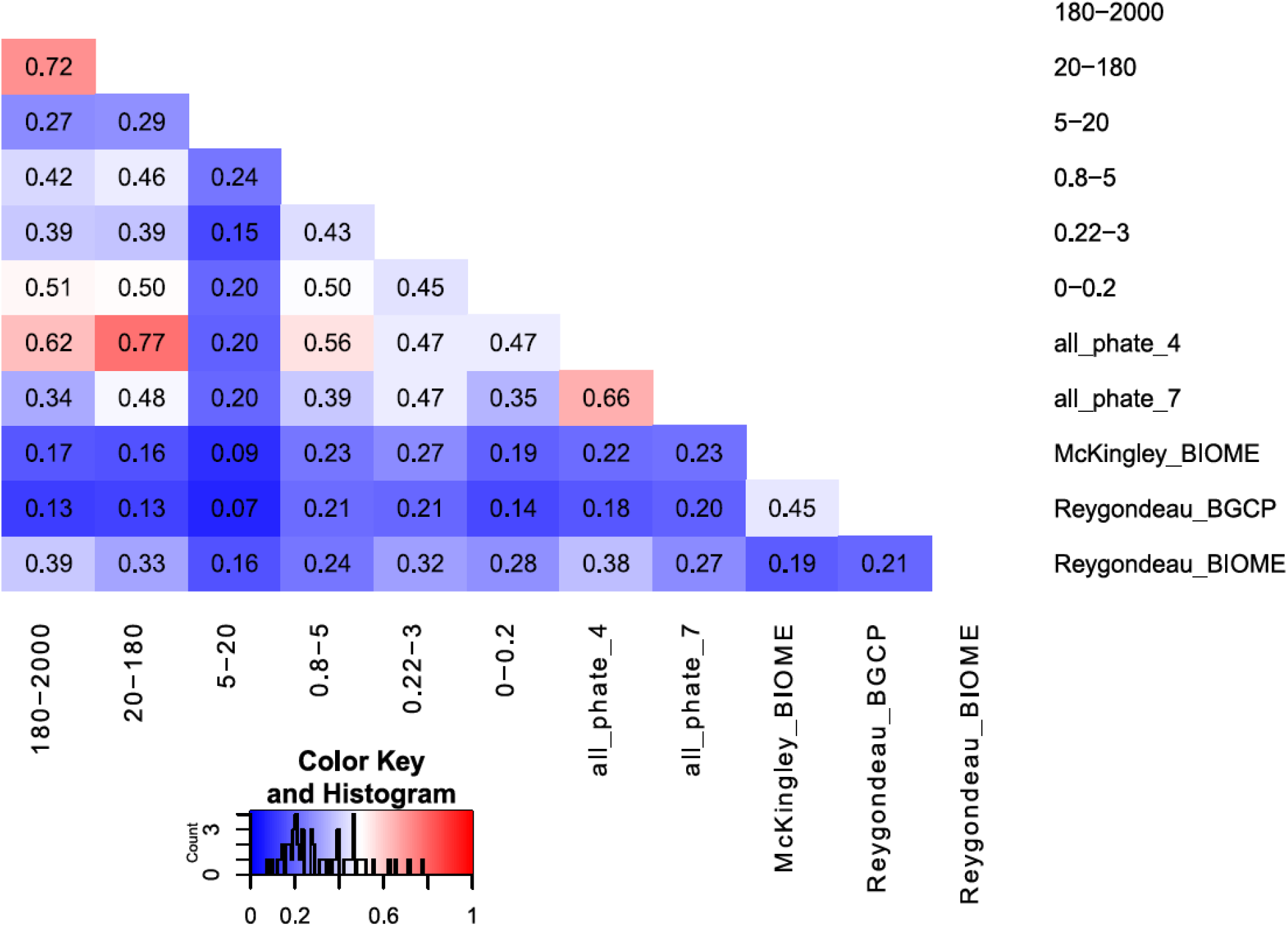
Pairwise comparisons of ocean partitions based on plankton provinces and existing biogeochemical partitions. The three oceanic partitions are Reygondeau et al.^10^ Biogeochemical Provinces (BGCP); Reygondeau et al. Biomes^10^; Fay and McKingley Biomes^11^.

**Supplementary Fig. 9.**
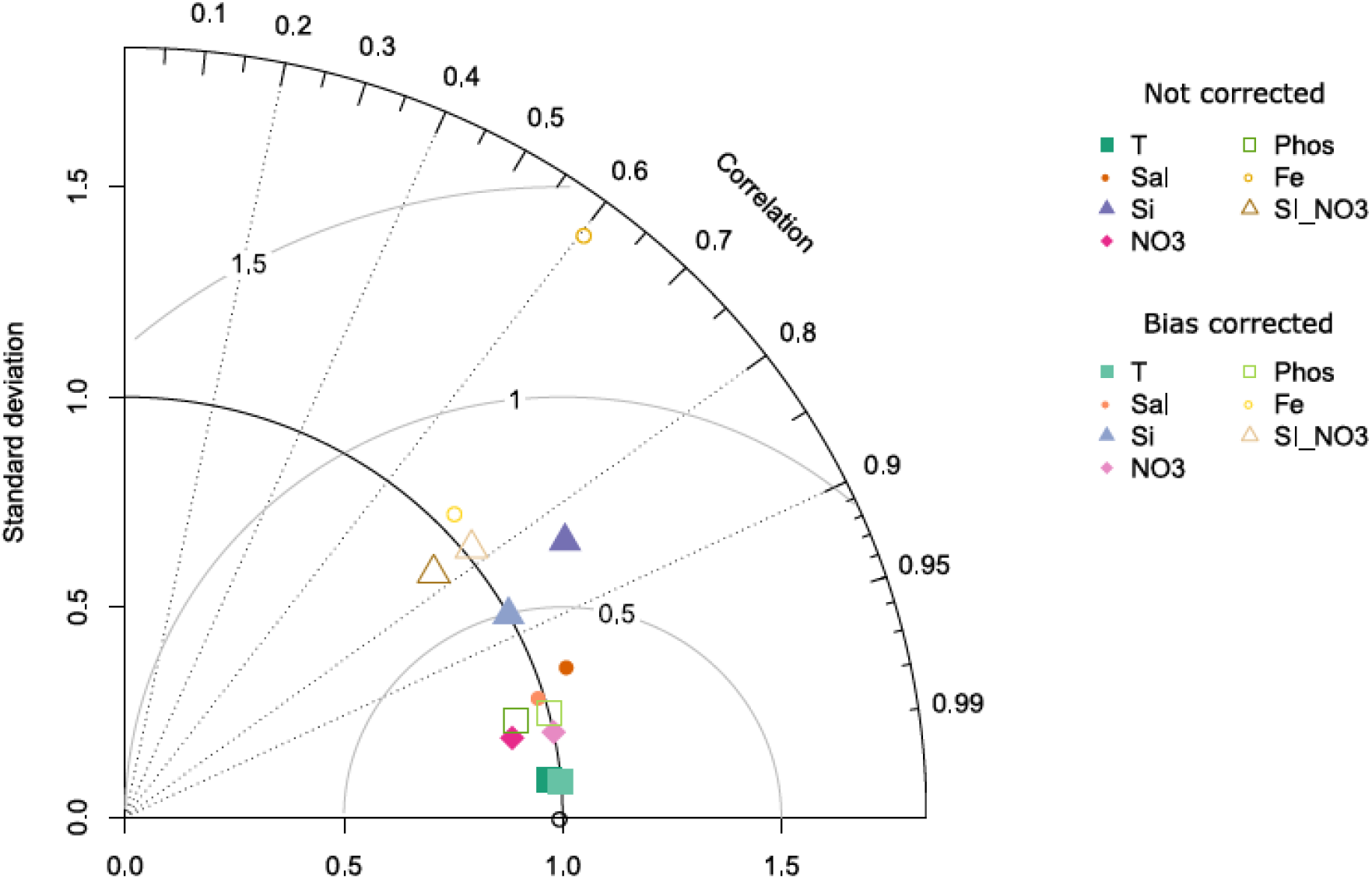
Taylor diagram exhibiting statistical comparison of WOA 2013 observations and present day ESM model mean of the different drivers. Each variable is centered and scaled according to the mean and standard deviation of the observed variable (black circle point at standard deviation 1 on the x-axis). Dark color points are the ESM model mean without Cumulative Distribution Function transform (CDFt^25^) bias correction. They are to be compared with light color points for which the bias correction is performed. Overall, good spatial correlations are found between the model mean and the observations. CDFt bias correction performs well by bringing standard deviations of the model to the observed standard deviations without decreasing correlations.

**Supplementary Fig. 10.**
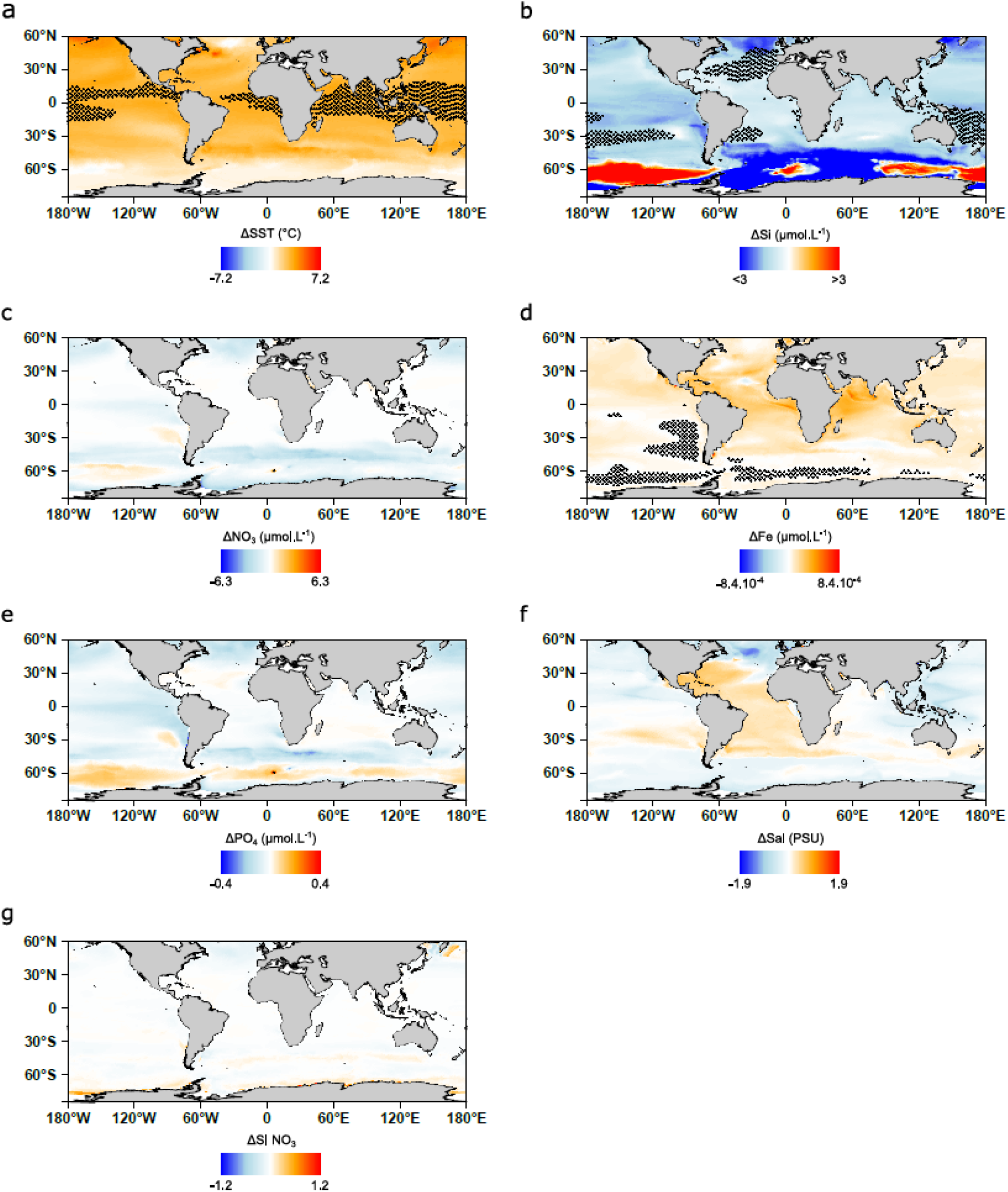
Differences in drivers’ intensity (2090/99-2006/15) in the bias corrected ESM model-mean under RCP8.5. (**a**) Sea Surface Temperature (SST). (**b**) Dissolved silica. (**c**) Nitrate. (**d**) Iron. (**e**) Phosphate. (**f**) Salinity (Sal). (**g**) Seasonality Index of Nitrate (SI NO_3_). Note that the scale for dissolved silica variations is restricted to visualize small variations. Regions for which out of range values (*i.e.* inferior to the minimum or superior to the maximum found in 2006-15) are reached at the end of the century are highlighted with small stars reflecting high uncertainty zones for machine learning approaches.

**Supplementary Fig. 11.**
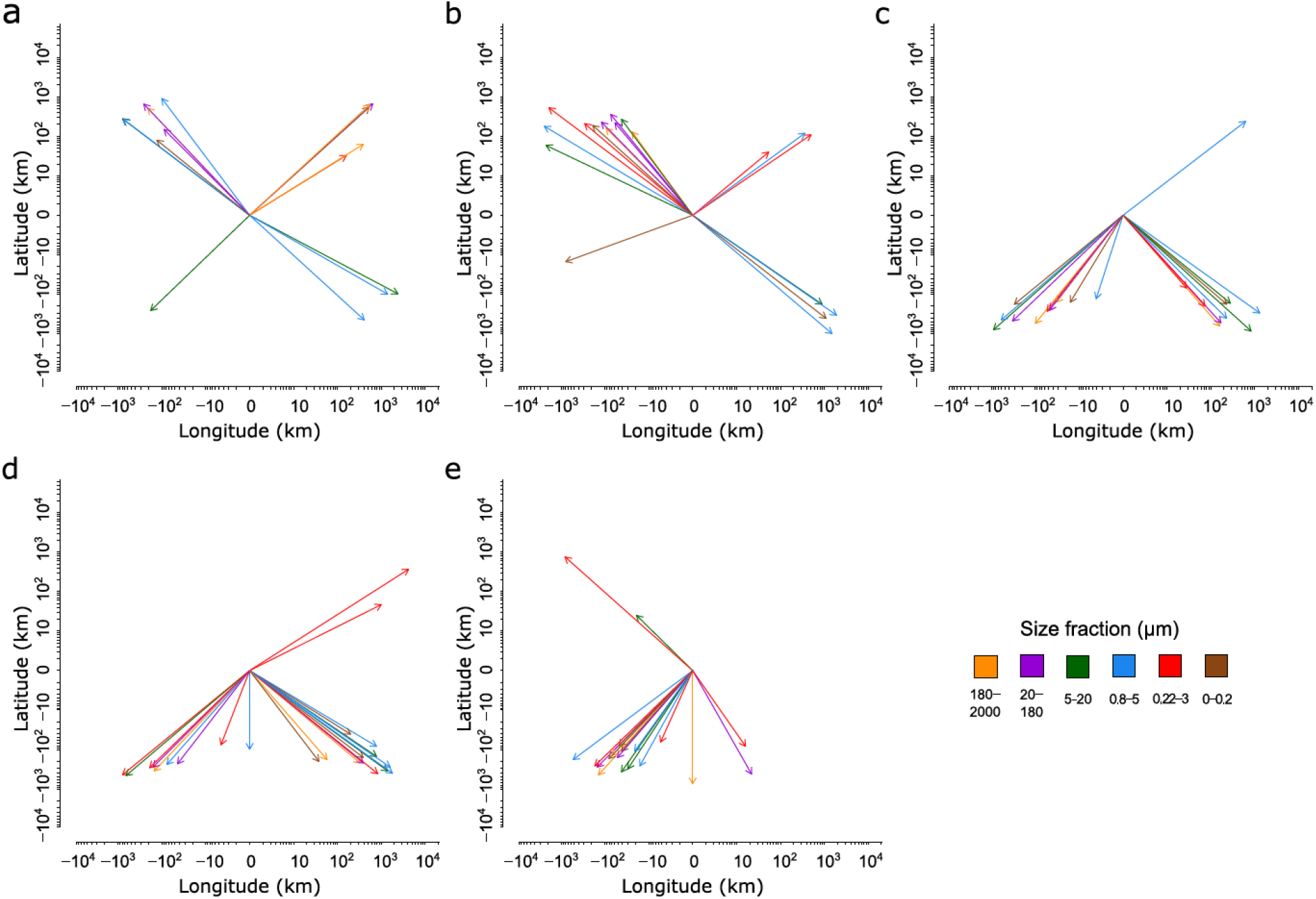
Projected migration shifts of the 27 provinces between present day and end of the century. Predicted migration shifts are presented in 5 major ocean basins: (**a**) North Atlantic (**b**) North Pacific (**c**) South Atlantic (**d**) South Pacific and (**e**) Indian Ocean. 96% of migration shifts (larger than 200 km) are oriented towards the pole. Mean shift is 641 km (76 ± 79 km.dec^-1^) and median shift is 394 km (47 km.dec^-1^). Few provinces are projected to shift more than thousands of kilometers towards suitable environmental conditions with a maximum shift of 4325 km.

**Supplementary Fig. 12.**
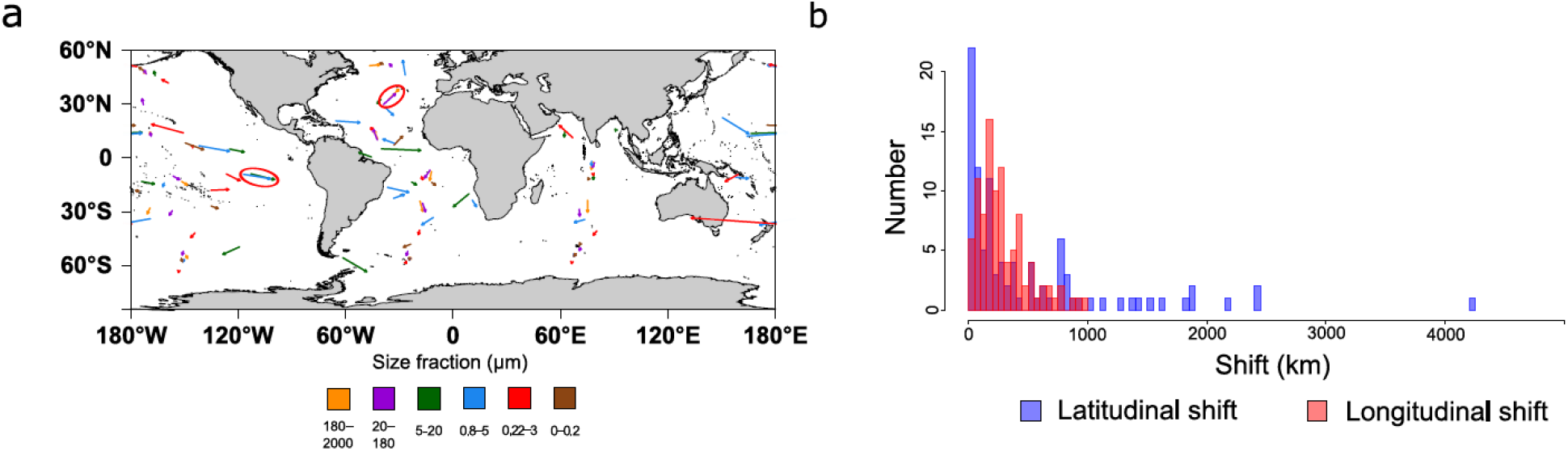
(a) Projected migration shifts of provinces on the world map. (b) Latitudinal shift distribution (red bars) and longitudinal shift distribution (blue bars). (**a**) Migration shifts are represented as arrows pointing at the end of century centroid. Arrows are colored according to the size fraction. Some shifts seem to correlate with each other (exemplified with circled arrows). For instance, parallel shifts are projected in the southern pacific equatorial communities of size fractions 0.8-5 µm and 5-20 µm (blue and green circled arrows). All non-poleward arrows belong to small size classes (<20µm) showing differential responses to climate change depending on the size class. (**b**) Some longitudinal shifts are more important than latitudinal shifts with 14 longitudinal shifts superior to 1000 kms. Mean longitudinal shift (around 500 kms) is significantly higher (Student t-test^26^ p-value<0.01) than mean latitudinal shift (around 290 kms) while medians (longitudinal 190 kms vs latitudinal 230 kms) are not significantly different (Wilcoxon test). The median migration speed is 47 km.dec^-1^, (latitudinally 23 km.dec^-1^, longitudinally 27 km.dec^-1^).

**Supplementary Fig. 13.**
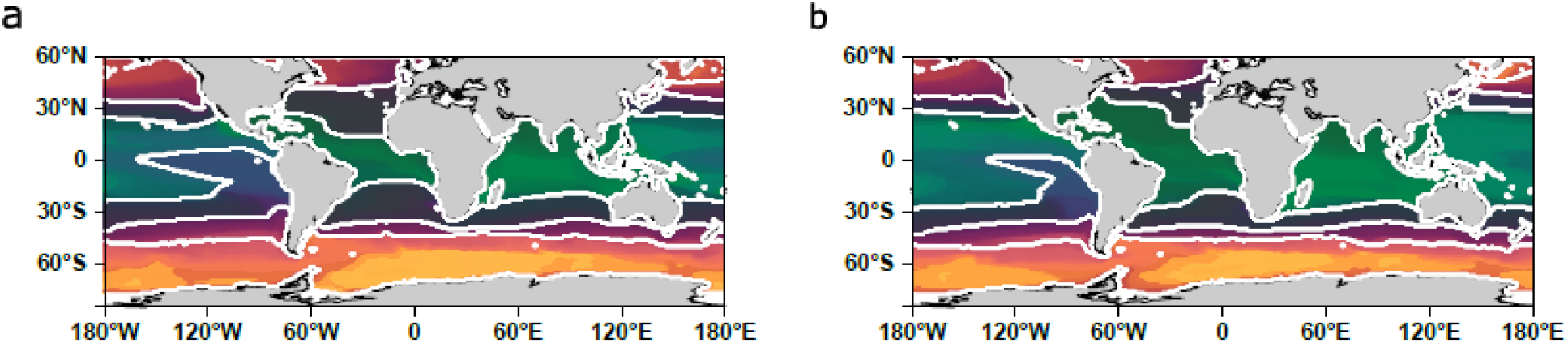
(a) Present day and (b) end of century combined size class biogeographies using the PHATE algorithm^7^. Each grid point is associated with a color depending on its PHATE coordinates (using three axes). Each coordinate is respectively assigned to a given degree of color between 0 and 255 according to equation 2 (either red green or blue depending on the axis). Boundaries of clusters using k-medoïds^27^ (k=4) are represented in white.

**Supplementary Fig. 14.**
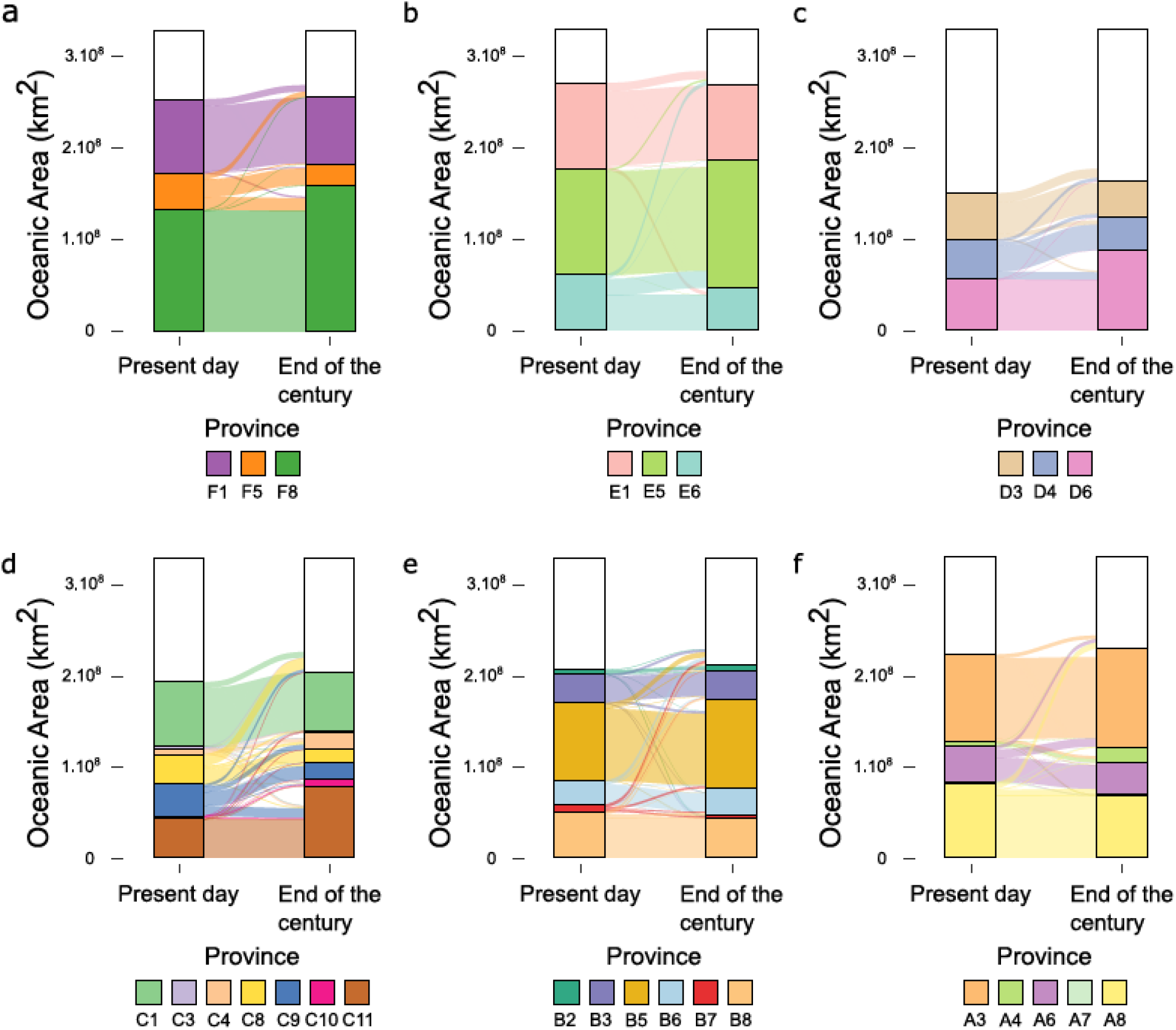
Probabilistic covered areas of the provinces projected in present day and at the end of the century. (**a**) 180-2000 μm (**b**) 20-180 μm (**c**) 5-20 μm (**d**) 0.8-5 μm (**e**) 0.22-3 μm (**f**) 0-0.2 μm. The area covered by a province is defined as the area in which this province is dominant and weighted by its probability of presence at each point and grid cell area. Areas not covered by the provinces are represented in white.

**Supplementary Fig. 15.**
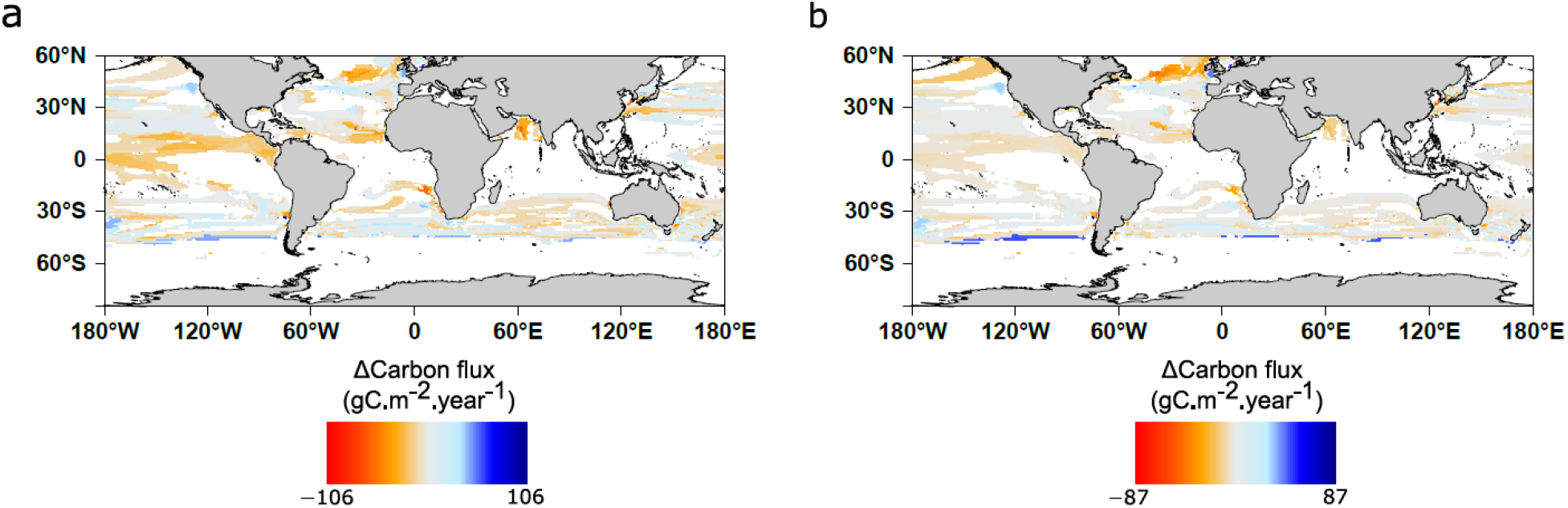
Differences in carbon export by the end of the century using data from two present day extrapolative models of carbon export. Each assemblage of *climato-genomic* province is associated to its mean export flux from present day extrapolative models. Then the difference between present and future export is calculated by the difference between export fluxes from future and present assemblages at a given point. Means for assemblages are calculated based on **(a)** Eppley et al.^28^ **(b)** Laws et al.^29^

**Supplementary Fig. 16.**
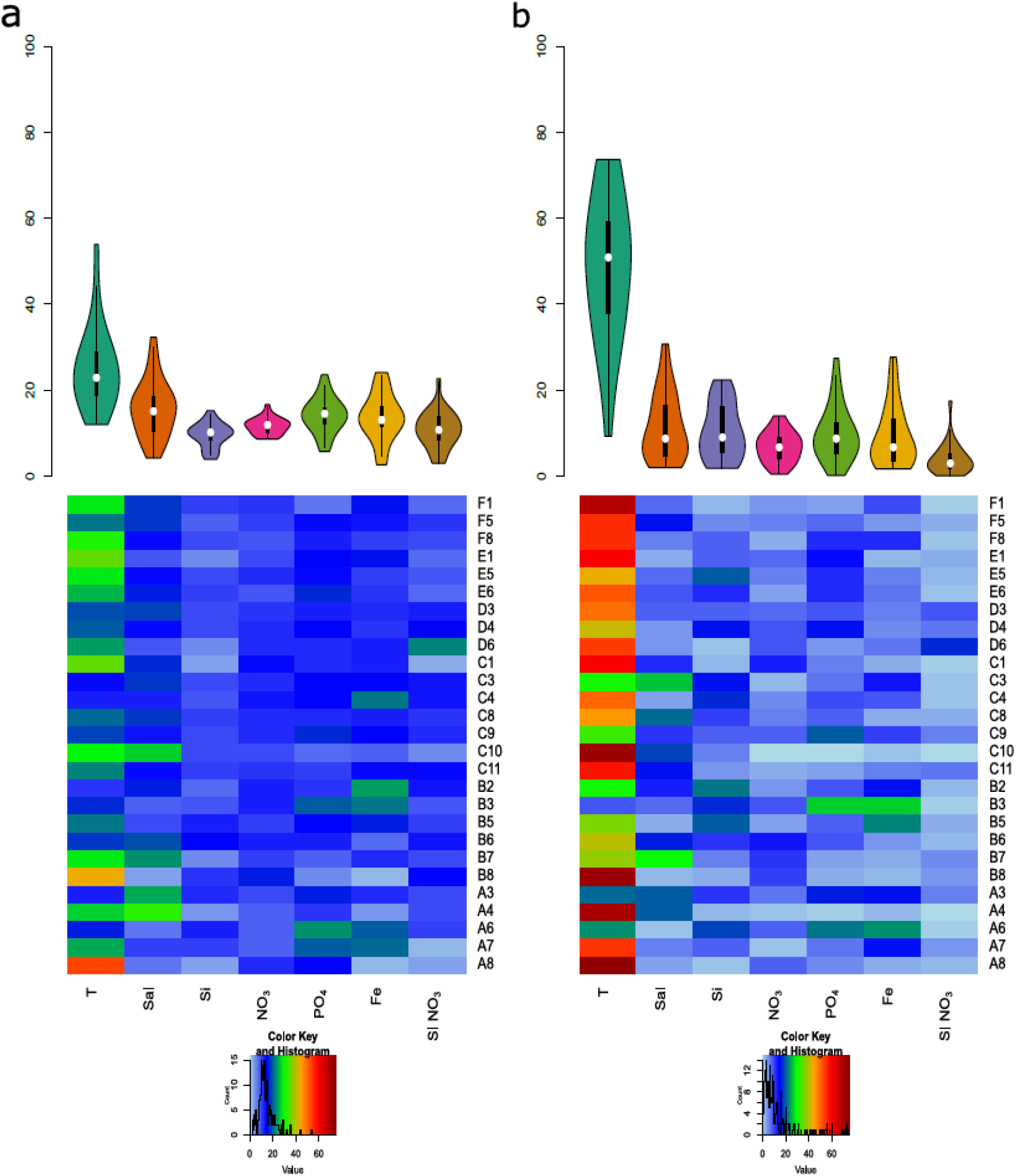
Distributions and heat maps of the relative influences of the different drivers in (a) defining single niches associated with the provinces from DALEX R package^30^ (b) driving climate change associated reorganization of single provinces. Median relative influence of Temperature is significantly higher than for all other environmental parameters (Pairwise Wilcoxon test p<0.01 for all parameters) in (**a**) defining the niches and (**b**) driving province reorganization. This is also the case within individual size fractions for (**b**) but not for (**a**). Respectively, Salinity (Sal) and Phosphate (PO_4_), have second and third highest median relative influences far behind Temperature whereas dissolved Silica and seasonality index of Nitrate (SI NO_3_) have the lowest median relative importance in (**a**). Numbers above violins are median (and mean) relative influences.

**Supplementary Fig. 17.**
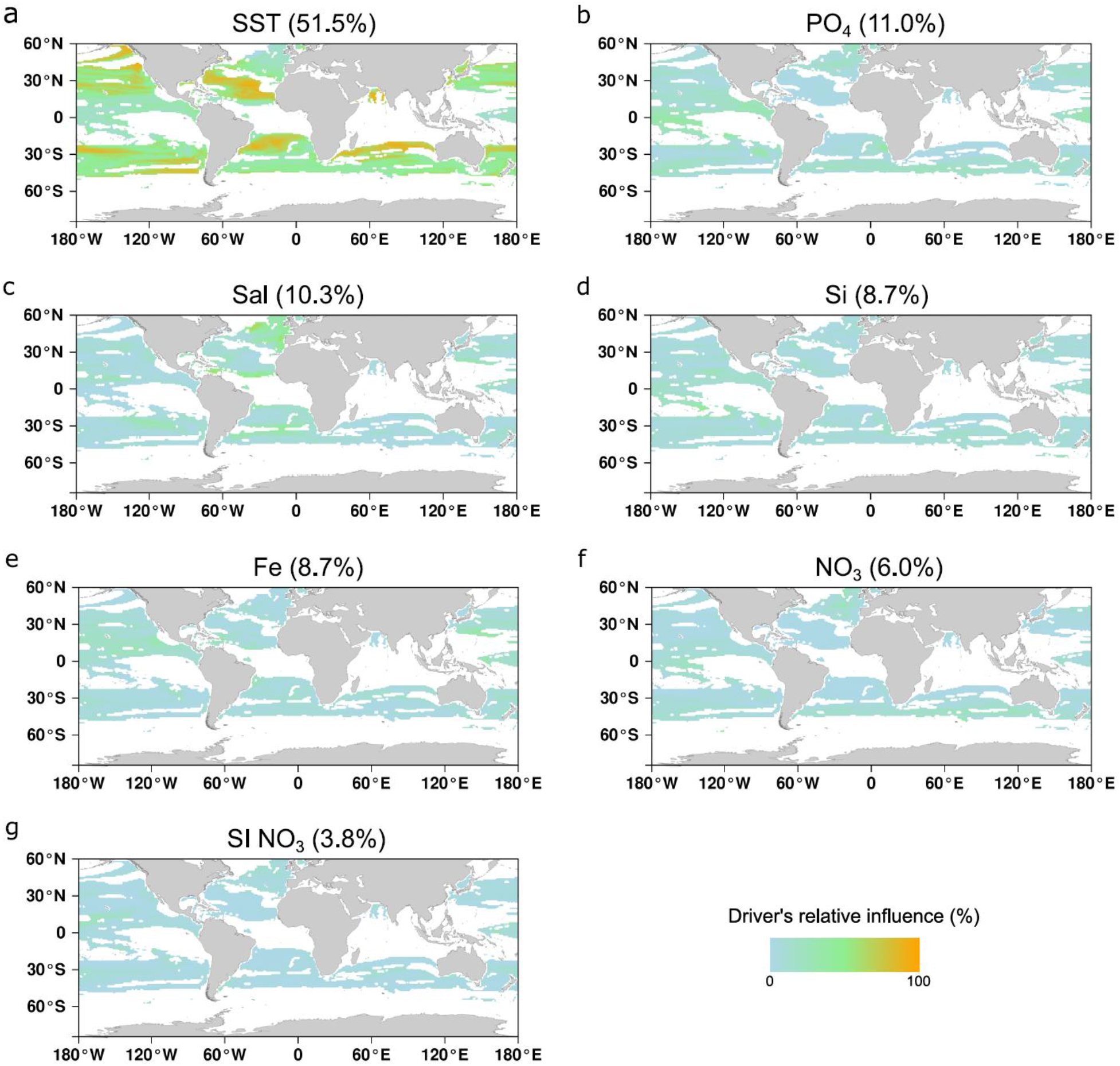
Maps of environmental drivers’ relative influences in driving province reorganization. Each environmental parameter’s relative influence is quantified by considering only the variation of each parameter individually between present day and end of the century as defined in Barton et al.^12^(*Materials and Methods*) and where a significant change is projected. Importantly the mean impact of SST (51.5%) is to a great extent the highest. PO_4_ is the second most impacting driver (11.0%). Contrary to Supplementary Fig. 18, relative influence is calculated here by combining all provinces together. Therefore, mean relative influences slightly differ especially for dissolved silica (Si).

**Supplementary Fig. 18.**
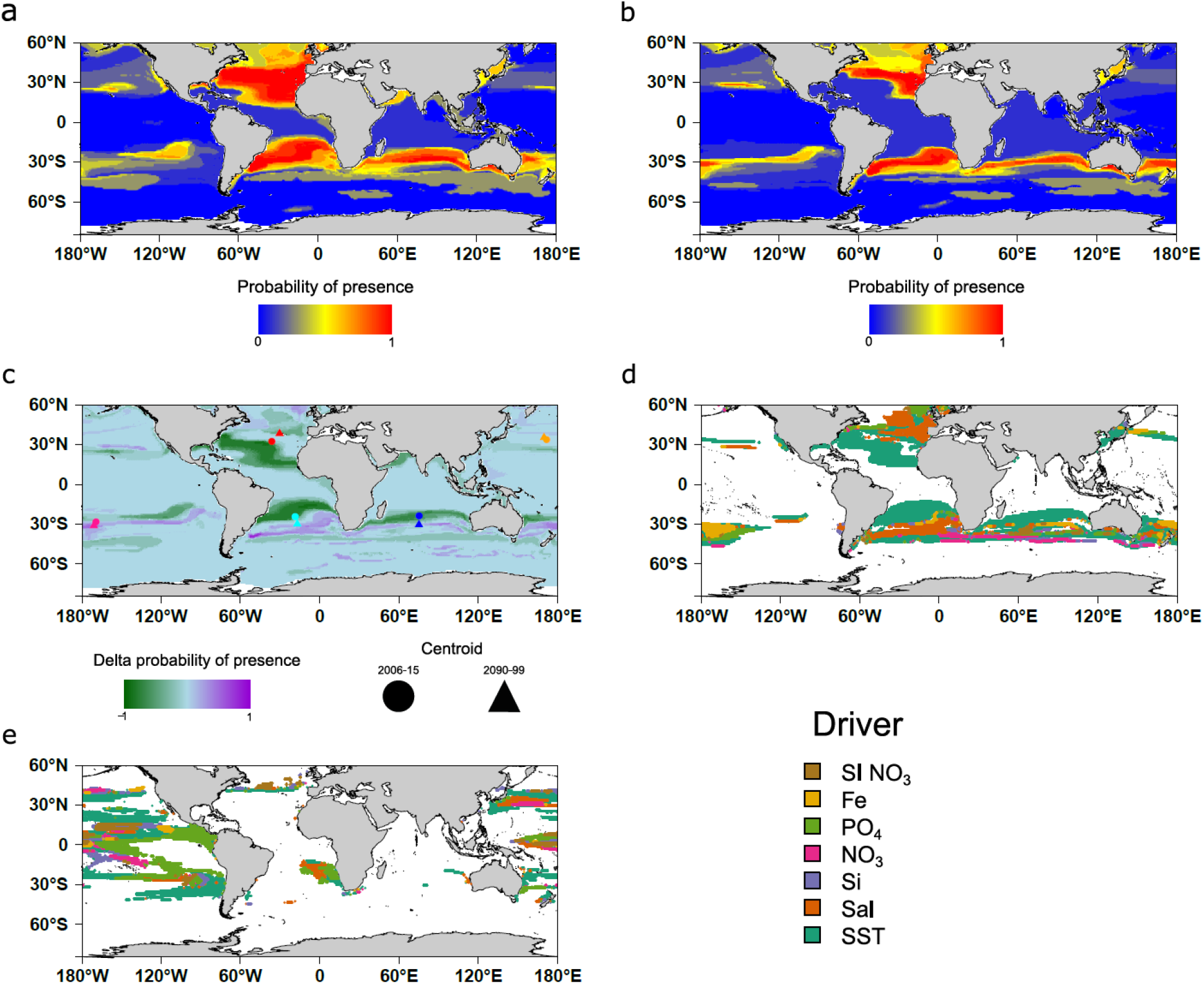
Projection maps of province F5 of size fraction 180-2000 μm in present day (a), at the end of the century (b) and their difference (c); Drivers of changes for province F5 (d) and C9 (e). At each grid point, the probability of presence of the province is computed as the average of the predicted probability of each of the four machine learning techniques (gbm, nn, rf and gam). Red color indicates a high probability of presence. (**a**)The projected province corresponds to the sampled province (North Atlantic and South Atlantic) but several other places have high probabilities of presence such as South Australia where no sampling is available. (**b**) At the end of the century, the province is projected to reduce significantly in size. (**c**) Delta probability of presence map (2006/15 – 2090/99) and core range shift in the 5 major oceanic basins of province F5 of size fraction 180-2000 µm. In all the basins, the centroid of the province is projected to migrate poleward. (**d**) Main drivers associated with the projected changes. Changes are mainly driven by sea surface temperature (56%) followed by salinity (16%). (**e**) Main drivers associated with the shrinkage of the equatorial cluster C9 of size fraction 0.8-5. Considering only latitudes between the two tropics, changes are mainly driven by decreases in PO_4_ (24%) in addition to SST (27%) (overall 34% STT and 20% PO_4_).

**Supplementary Table 1.** Earth System models used to compute the mean model.

| Model | Reference |
| --- | --- |
| CESM1-BGC | Gent et al., 2011<sup>31</sup> |
| GFDL-ESM2G | Dunne et al., 2013<sup>32</sup> |
| GFDL-ESM2M | Dunne et al., 2013<sup>32</sup> |
| HadGEM2-ES | Collins et al., 2011<sup>33</sup> |
| IPSL-CM5A-LR | Dufresne et al., 2013<sup>34</sup> |
| IPSL-CM5A-MR | Dufresne et al., 2013<sup>34</sup> |
| MPI-ESM-LR | Giorgetta et al., 2013<sup>35</sup> |
| MPI-ESM-MR | Giorgetta et al., 2013<sup>35</sup> |
| NorESM1-ME | Bentsen et al., 2013<sup>36</sup> |

**Supplementary Table 2.** Genomic provinces climatic annotations and oceanic surfaces covered in present day (2006-15) and at the end of the century (2090-990).

| Fraction | Province | Climatic annotation | Area 2006-15<br>(MKm<sup>2</sup>) | Area 2090-99<br>(MKm<sup>2</sup>) | Delta area<br>(MKm<sup>2</sup> (%)) |
| --- | --- | --- | --- | --- | --- |
| 180-2000 | F1 | polar | 82 | 73 | -9 (-11%) |
| 180-2000 | F5 | temperate | 44 | 26 | -18 (-41%) |
| 180-2000 | F8 | tropico-equatorial | 140 | 169 | +29 (+21%) |
| 20-180 | E1 | polar | 97 | 84 | -13 (-13%) |
| 20-180 | E5 | tropico-equatorial | 119 | 145 | +26 (+22%) |
| 20-180 | E6 | temperate | 65 | 49 | -15 (-23%) |
| 5-20 | D3 | temperate | 51 | 41 | -10 (-20%) |
| 5-20 | D4 | equatorial | 46 | 37 | -8 (-18%) |
| 5-20 | D6 | tropical | 65 | 97 | +32 (+48%) |
| 0.8-5 | C1 | polar | 71 | 64 | -6 (-8%) |
| 0.8-5 | C3 | subtropical | 3,6 | 1,4 | -2,2 (-61%) |
| 0.8-5 | C4 | subtropical | 6 | 19 | +13 (+217%) |
| 0.8-5 | C8 | temperate | 33 | 15 | -18 (-54%) |
| 0.8-5 | C9 | equatorial | 35 | 19 | -16 (-45%) |
| 0.8-5 | C10 | subtropical | 3.8 | 7.4 | +3,6 (+95%) |
| 0.8-5 | C11 | tropical | 49 | 86 | +37 (+75%) |
| 0.22-3 | B2 | subtropical | 5.1 | 6.7 | +1,6 (+31%) |
| 0.22-3 | B3 | equatorial | 33 | 32 | -1 (-0.3%) |
| 0.22-3 | B5 | tropical | 88 | 102 | +13 (+15%) |
| 0.22-3 | B6 | temperate | 30 | 35 | +5 (+17%) |
| 0.22-3 | B7 | temperate | 8 | 2 | -6 (-75%) |
| 0.22-3 | B8 | polar | 51 | 44 | -7 (-14%) |
| 0-0.2 | A3 | tropical | 101 | 114 | +13 (+13%) |
| 0-0.2 | A4 | subtropical | 6 | 19 | +13 (+216%) |
| 0-0.2 | A6 | equatorial | 40 | 36 | -4 (-10%) |
| 0-0.2 | A7 | temperate | 1.7 | 1 | -0.7 (-41%) |
| 0-0.2 | A8 | polar | 81 | 67 | -14 (-17%) |

**Supplementary Table 3.**
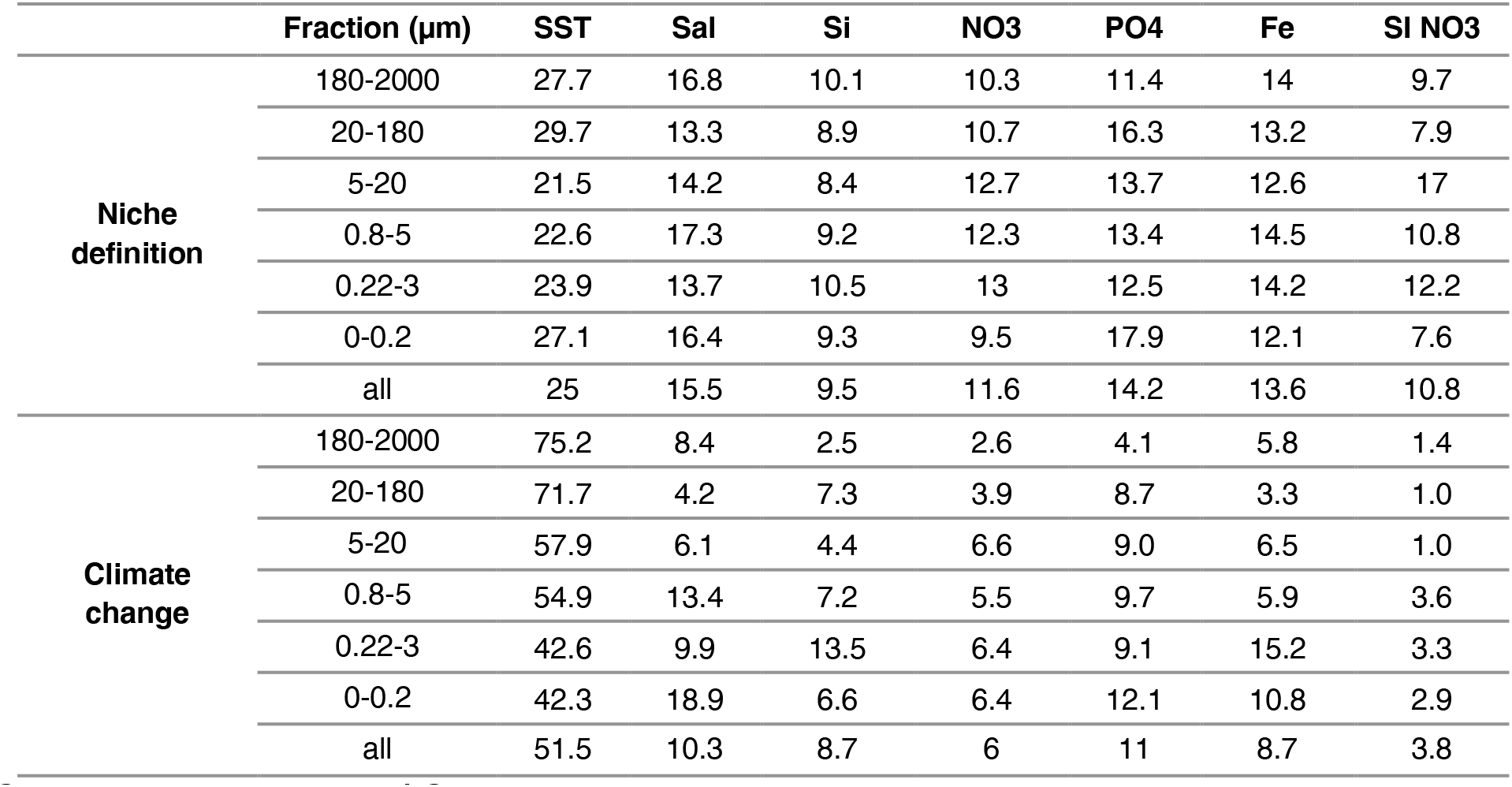
Summary table of the relative importance of each environmental driver in niche definition and in driving geographical reorganization in response to climate change. Note that in both cases (niche definition and climate change), the row ‘all’ is not the mean over the size fractions. In the case of niche definition, this is due to a different number of niches in each size class. In the case of climate change, relative influence is either calculated for single provinces at a given grid point then recalculated for individual size class or calculated for all provinces together (row ‘all’).

| | Fraction ($\mu\text{m}$) | SST | Sal | Si | NO3 | PO4 | Fe | SI NO3 |
| --- | --- | --- | --- | --- | --- | --- | --- | --- |
| **Niche definition** | 180-2000 | 27.7 | 16.8 | 10.1 | 10.3 | 11.4 | 14 | 9.7 |
|  | 20-180 | 29.7 | 13.3 | 8.9 | 10.7 | 16.3 | 13.2 | 7.9 |
|  | 5-20 | 21.5 | 14.2 | 8.4 | 12.7 | 13.7 | 12.6 | 17 |
|  | 0.8-5 | 22.6 | 17.3 | 9.2 | 12.3 | 13.4 | 14.5 | 10.8 |
|  | 0.22-3 | 23.9 | 13.7 | 10.5 | 13 | 12.5 | 14.2 | 12.2 |
|  | 0-0.2 | 27.1 | 16.4 | 9.3 | 9.5 | 17.9 | 12.1 | 7.6 |
|  | all | 25 | 15.5 | 9.5 | 11.6 | 14.2 | 13.6 | 10.8 |
| **Climate change** | 180-2000 | 75.2 | 8.4 | 2.5 | 2.6 | 4.1 | 5.8 | 1.4 |
|  | 20-180 | 71.7 | 4.2 | 7.3 | 3.9 | 8.7 | 3.3 | 1.0 |
|  | 5-20 | 57.9 | 6.1 | 4.4 | 6.6 | 9.0 | 6.5 | 1.0 |
|  | 0.8-5 | 54.9 | 13.4 | 7.2 | 5.5 | 9.7 | 5.9 | 3.6 |
|  | 0.22-3 | 42.6 | 9.9 | 13.5 | 6.4 | 9.1 | 15.2 | 3.3 |
|  | 0-0.2 | 42.3 | 18.9 | 6.6 | 6.4 | 12.1 | 10.8 | 2.9 |
|  | all | 51.5 | 10.3 | 8.7 | 6 | 11 | 8.7 | 3.8 |

## Notes

### Competing Interest Statement

The authors have declared no competing interest.

